# A comprehensive pharmacodynamic dataset reconciles 50 years of laboratory and clinical knowledge

**DOI:** 10.1101/2025.09.22.676882

**Authors:** Eugene F Douglass, Chinmay Joshi, Emily Hannan, Tobin Paez, Elizabeth A. Hughes, Jonathan P. Mochel

**Affiliations:** Department of Pharmaceutical and Biomedical Sciences, University of Georgia, 250 W. Green Street, Athens, GA 30602; Precision One Health Initiative, University of Georgia College of Veterinary Medicine, Athens, GA, USA

## Abstract

Single-agent clinical response data for widely used chemotherapies have remained difficult to analyze because they are scattered across decades of print literature. We consolidated these sources into NCI1970-Meta, a 49,002-patient dataset covering 30 drugs across 18 cancers, enabling the first quantitative comparison of clinical outcomes, laboratory potency metrics, clinical exposure, and literature-derived biomarkers.

Clinical patterns were strongly lineage-driven: cancer type explained far more ORR variability than drug identity (37.3% vs 15.6%; F = 13.43 vs 3.29). FDA approvals reflected these same patterns where ORR strongly predicted indication status (F = 98.3-104.1).

In contrast, laboratory efficacy metrics did not track clinical activity. Raw in vitro AUC showed no association with ORR (R2 = 0.00; 95% CI: 0.00-0.01) and was dominated by drug identity rather than cancer lineage (85.2% vs 8.6%; F = 212.2 vs 21.6). Instead, AUC correlated with clinical exposure: unbound Cmax (R2 = 0.21 [0.21-0.34]) and therapeutic minimum concentrations (R2 = 0.69 [0.64-0.73]). This indicates that standard assay ranges capture exposure requirements rather than true efficacy. Normalizing potency by exposure restored the expected clinical relationships and resolved drug-specific anomalies such as gemcitabine.

Biomarkers showed consistent behavior across clinical and laboratory settings. Among 314 biomarker-drug pairs, correlation directions were significantly conserved (R2 = 0.17 [0.10-0.25]; p = 2.8×10^−14^). Literature-defined sensitivity and resistance annotations were enriched in vitro (OR = 3.7; p = 2.36×10^−8^) and in the clinic (OR = 3.9; p = 6.52×10^−9^), with stronger performance for correlations >0.1 (OR = 13.3 in vitro; OR = 8.4 clinically). Simple biomarker-sum models performed well across drugs, consistent with multi-pathway pseudo-first-order behavior.

Overall, NCI1970-Meta provides a quantitative framework linking laboratory pharmacology to real-world clinical efficacy. Biological signal is reliably preserved within drugs, while cross-drug comparisons require explicit exposure normalization. This resource offers a statistical foundation for improving drug prioritization, biomarker development, and translational pharmacodynamic modeling.

## INTRODUCTION

### Clinical-Laboratory “efficacy” dichotomy

High-throughput drug screening has generated millions of *in vitro* dose-response curves across hundreds of cancer cell lines and FDA-approved agents.^1-4^ These assays typically define activity by the concentration that reduces cell viability by 50% (IC50), or area under the curve (AUC) creating a potency-based view of drug action (Fig. 1A). In contrast, clinical oncology defines efficacy by tumor shrinkage over time, measured as overall response rate (ORR) during Phase 2-3 trials (Fig. 1B).^5^ The disconnect between laboratory dose-response metrics and clinical time-response outcomes has limited the translational utility of preclinical assays (Fig. 1C).^6^

**Figure 1.**
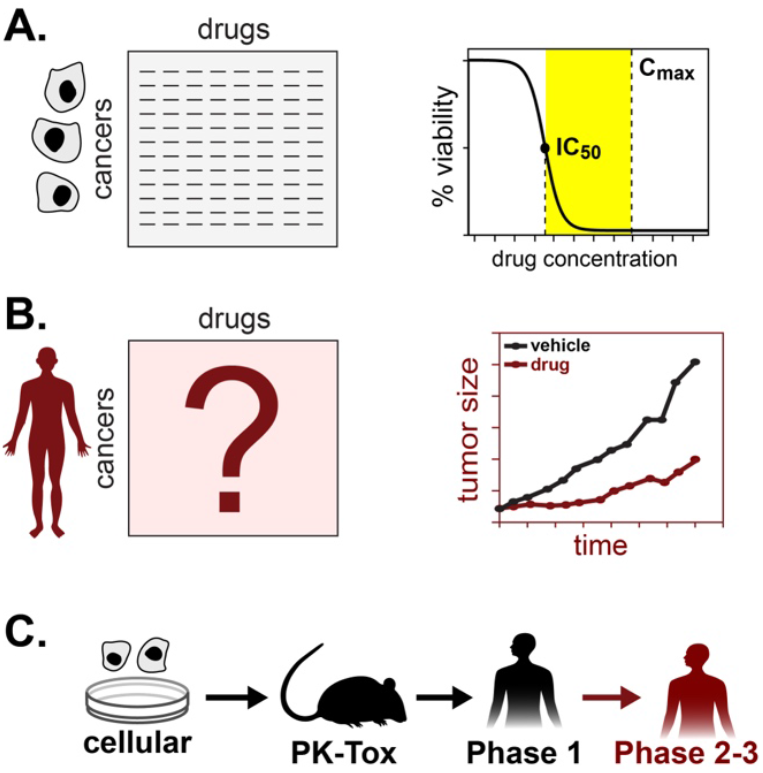
Data-gap (Clinical response rates) filled by this study. (A) Most *in vitro* drug response data is defined by *dose-response metrics* such as AUC or IC50, (B) In contrast, clinical response data is defined by *time-response metrics* which is currently not available in a digitized form. (C) Drug development proceeds from cellular assays to animal PK-toxicity, Phase 1 dose-escalation, and finally Phase 2-3 efficacy trials. We align summary statistics across these stages by comparing *in vitro* IC50 values with, Phase 1 drug-exposure and clinical response rates.

### Concentration-response metrics

This gap also appears in how preclinical potency metrics are used during early drug development. In vitro IC50 values, measured in both disease and normal models, are commonly treated as proxies for the therapeutic minimum (TRmin) and therapeutic maximum (TRmax), the concentration boundaries later formalized during Phase 1 dose-escalation studies.^7,8^ At the same time, these same IC50/AUC values are interpreted as measures of drug-responsiveness,^2-4^ raising a critical question: how much in vitro metrics inform *clinical efficacy (ORR)* compared with the clinical *concentration range(TR)*:

- *IC*_50_/*AUC* ∝ ***TR***?
- *IC*_50_/*AUC* ∝ ***ORR*** ?

This distinction matters clinically, because diagnostic biomarker studies and drug-repurposing efforts often begin with simple concentration-response measurements in cell-line patient-derived samples.^9-11^

A practical way to address this question has been to normalize in vitro drug-potency (IC50) by clinically achievable exposure using a “clinical relevance ratio” (CRR) defined as:^12^

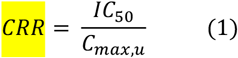

Where Cmax,u is the maximum achievable unbound plasma concentration of a drug under standard regimens.^13^ This metric evaluates whether an observed in vitro effect occurs at concentrations that patients can realistically reach. CRR < 1 suggest that efficacy or toxicity is achievable at clinical exposure levels, where as CRRs > 1 indicate that either outcome is unlikely at physiologic concentrations. This approach has been formally recommended the FDA for interpreting in vitro drug-drug interactions and toxicity studies,^12^ adopted to improve drug-repurposing pipelines for SARS-CoV-2^14^ and is now widely used in patient-derived organoid assays^15-18^ to assess whether dose-response effects observed ex vivo are consistent with concentrations attainable in patients.

### Time-response metrics

Although the CRR framework reconciles *drug-potency* measurements, it does not resolve the deeper mismatch between how *drug-efficacy* is defined in the lab and in patients. *In vitro* assays typically quantify activity using the area under the curve (AUC), which captures both dose-response steepness and maximal inhibition (Fig. 1A).^6^ Clinical evaluations, however, are time-centric: benefit is defined by reductions in disease burden over weeks to months. In oncology, RECIST criteria classify response by measurable decreases in tumor size and define non-response as continued growth (Fig. 1B).^5,19,20^ These differing endpoints underscore the need to understand how laboratory metrics such as AUC relate to clinical measures like overall response rate (Fig 1).^6^

Linking these stages remains challenging because clinical single-agent ORR data are fragmented, often buried in decades-old monographs or meta-reviews, and rarely digitized for systematic analysis. ^21^ Without this clinical anchor, efficacy values measured *in vitro* can easily give *false-negatives* when drugs that reach high plasma levels (e.g. of 5-fluorouracil or ifosfamide with Cmax,u is ∼430 µM) may appear inactive in laboratory screens capped at 5 µM.^1-4^

### Clinical single-agent data

This disconnect is compounded on the clinical side. Because most modern oncology trials evaluate drug combinations, contemporary single-agent efficacy data are scarce (Fig 2).^22^ The foundational evidence for many chemotherapies instead comes from national monotherapy trials conducted in the 1960s-1980s under the National Cancer Institute (NCI), which enrolled thousands of patients and directly measured single-agent response rates.^21^ Yet these results remained scattered across monographs, internal reports, and review articles, never systematically digitized or linked to laboratory metrics. As a result, the clinical endpoints needed to anchor laboratory potency particularly have been largely inaccessible to modern computational or biomarker-driven oncology. ^21^

**Figure 2.**
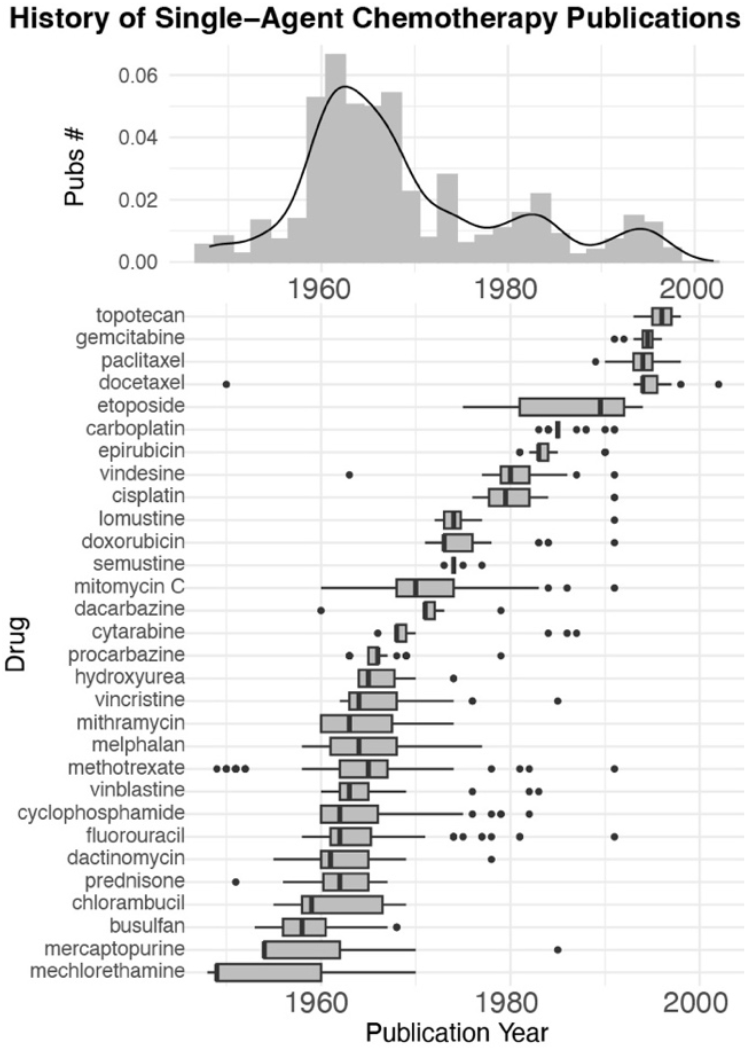
Historical Overview of Single-Agent Phase 2-3 studies on cytotoxic chemotherapies. Histogram illustrates the frequency of compiled single-agent publications from 1955-2000 which peaked in the 1960-70’s. Boxplot illustrates the time-periods over current cytotoxic chemotherapies were initially investigated.

To address this gap, we created *NCI1970meta*, a harmonized dataset of 49,002 patients across 931 single-agent chemotherapy trials, covering 30 drugs and 18 cancer types. By digitizing and reconciling these historical data^21^ we provide the first large-scale, quantitative map Phase 2-3 single-agent efficacy outcomes across 30 of the most important drugs in oncology (Fig 2).^23^ In addition, we have digitized additional pharmacodynamic and pharmacodynamic datasets that clarify efficacy trends including: Phase 1 pharmacodynamic boundaries (TRmin, TRmax), clinical exposure metrics (Cmax, AUC), and a comprehensive set of 234 biomarkers from the primary literature.^13,22,24-28^ Overall, this dataset allows us to systematically benchmark the relationship between laboratory potency metrics and clinical response, clarifying when and why translational failures occur.

### Laboratory single-agent data

To benchmark laboratory metrics against clinical outcomes, we used the three largest high-throughput drug-screening resources generated by the Broad Institute and the Sanger Institute: the Cancer Therapeutics Response Portal (CTRP), the Genomics of Drug Sensitivity in Cancer (GDSC), and the PRISM barcoded-pool platform.^2-4^ CTRP and GDSC each profile roughly 907 and 969 cancer cell lines with substantial overlap.^2,3^ They include broad representation of hematopoietic, epithelial, and mesenchymal cancers, which makes them essential for cancer-type analyses and for biomarker studies across lineages. However, their compound libraries are relatively small (297 or 545) and only contain a fraction of clinically approved agents.^2,3^ In contrast, PRISM focuses on a clinically relevant drug collection that includes 1,448 FDA-approved drugs and nearly complete coverage of approximately 143 commonly used oncology agents, but its cell-line library is limited to 497 adherent, mostly epithelial cancers.^4^ As a result, CTRP and GDSC are stronger for evaluating biological variability across cancer types,^2,3^ while PRISM is stronger for evaluating variability across drugs.^4^ All conclusions in this study were tested across all three datasets using sensitivity analyses, and the overall patterns were consistent: biological signal was strongest in GDSC and CTRP, and drug-level signal was strongest in PRISM (Fig S5-6).

### Biomarker analyses

Finally, we conducted a comprehensive evaluation of 234 literature-defined biomarkers across our full drug library to assess their performance in both clinical and in vitro settings.^27,28^ This analysis fills a long-standing gap that very few laboratory-defined biomarkers of chemotherapy response have been evaluated clinically.^29^ For example, in non-small cell lung cancer, where more than eighty cytotoxic chemotherapy biomarkers have been proposed, yet only thirteen have ever been evaluated in patients and only four show any diagnostic promise.^30^ Our results support the emerging consensus that single biomarkers are insufficient.^10,30^ Across both datasets, summed multi-gene biomarker sets consistently outperformed individual markers, particularly for anthracyclines, taxanes, and vinca alkaloids. These findings underscore the need for systematic clinical benchmarking to prioritize and integrate robust multi-marker signatures for cytotoxic chemotherapy.

#### Dataset Curation

### Initial monograph

Our study began with the NCI monograph *Single Agents in Cancer Chemotherapy*, which compiled national monotherapy trials conducted between 1955 and 1970.^21^ Although many trials were never formally published, the monograph reported evaluable patients and responder counts in a consistent format. We digitized these data and organized them into a drug-cancer matrix containing 23,222 patients across 463 trials, covering 17 drugs and 88 cancer labels that we consolidated into 26 TCGA-defined cancer types (Fig S1-3, Supp Data 1-2). From this raw resource, we focused on a clinically relevant subset of 19,958 evaluable patients spanning 30 drugs and 18 cancer types, including 14 WHO-defined essential cancer drugs not present in the original monograph.^23^ However, this dataset was sparse, with 58% of drug-cancer pairs missing data (Fig. S4A).

### Subsequent meta-reviews

To reduce sparsity, we integrated data from 48 meta-reviews published between 1970 and 2000 (Fig S2). These reviews summarized Phase II-III single-agent trials for specific drugs or cancers and often provided larger, more robust cohorts than the original NCI entries.^21^ To avoid duplication, we combined manual extraction of response numbers and bibliographic data from all recoverable primary sources with a rule-based algorithm that selected the most comprehensive dataset for each drug-cancer pair (Fig. S2-3). This two-pronged approach was necessary because many meta-reviews incorporated unpublished aggregation results that could only be included using our rule-based procedure. The final reconciled dataset, NCI1970meta, contains 49,002 evaluable patients treated with 30 chemotherapies across 18 malignancies. For each trial, both total patients and responders are recorded, enabling standardized calculation of overall response rate (ORR) across 540 drug-cancer combinations.^5,19,20^

## RESULTS

### Clinical vs Laboratory Metric Analyses

#### Clinical ORR mirrors FDA approval

We next compared clinical response rates from NCI1970-Meta with FDA-indications and *in vitro* screening datasets to assess how well laboratory efficacy metrics reflect real clinical activity (Fig. 3). Overall response rates (ORRs) showed a striking and consistent pattern: hematologic malignancies had the highest responses, whereas gastrointestinal cancers were among the least responsive (Fig. 2A-B). This pattern was supported by two-way ANOVA, which showed that cancer type accounted for far more ORR variance than drug identity (37.3% vs 15.6%; F = 13.43 vs 3.29).

**Figure 3.**
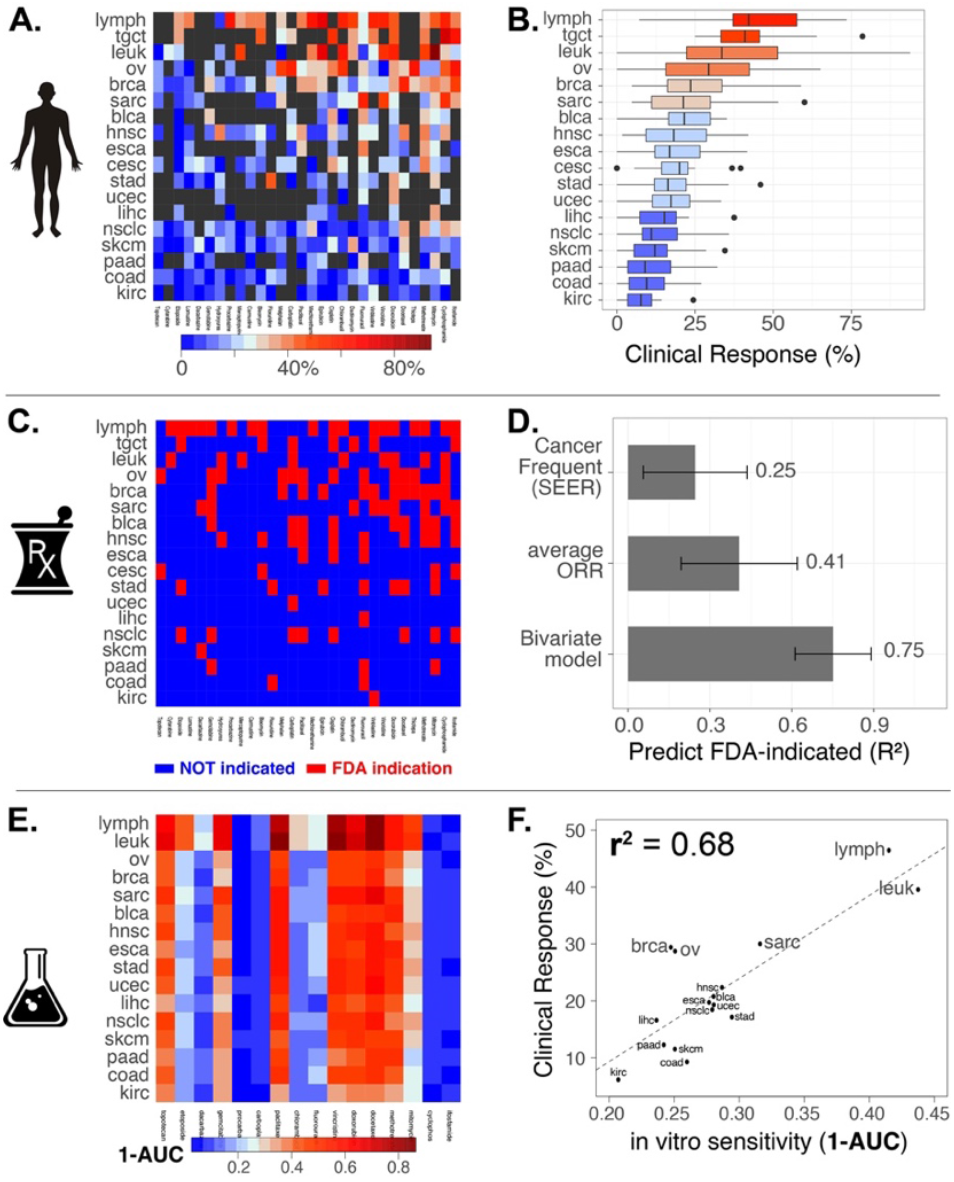
“Rosetta stone” analysis: NCI1970meta, FDA-indications and laboratory-efficacy matrices are plotted as heatmaps with the same rank ordering of cancer-types and drugs. **(A)**. Heatmap of ORR data with drugs and cancers sorted by highest average ORR **(B)** Boxplots of ORR across cancer-types **(C)** FDA-indications^22^ visually mirror NCI1970-meta ORR data **(D)** Multi-variate analysis reveals that the number of FDA-indications appears to be a function of cancer chemosensitivity and frequency of the cancer in the USA. **(E)**. *In vitro* drug-sensitivity appears visually distinct from both FDA-indication and ORR data **(F)** The lineage-dependence of drug-response is conserved in *in vitro* and in the clinic.

The same lineage-level hierarchy appeared in the current landscape of FDA single-agent approvals (Fig. 2C),^22^ and this association was statistically conserved (F = 5.6 vs 2.2). Weighted and unweighted ordinary least squares models confirmed that ORR strongly predicted FDA-indication status (F = 98.3 and 104.1, respectively). Moreover, multivariate analysis showed that both ORR and the prevalence of each cancer type in the United States independently predicted FDA-indication status (R^2^ = 0.75, Fig. 2D),^31^ indicating that FDA-approvals reflect both efficacy and the logistical feasibility of enrolling sufficient patients.

#### In vitro efficacy metrics fail to mirror clinical activity

In contrast to the strong clinical patterns described above, raw in vitro drug efficacy metrics such as (1-AUC) did not resemble either clinical response rates or FDA-approval patterns (Fig 3E).^2,3^ Variance decomposition analysis revealed that drug identity overwhelmingly dominated AUC variability compared with cancer type (85.2% vs 8.6%; F = 212.2 vs 21.6), indicating that class-specific chemical properties overshadow lineage-specific biology. Visually, this appears as vertical streaks in the in vitro heatmaps, in sharp contrast to the horizontal clinical patterns driven by cancer type (Fig 3 “Rosetta Stone”). Notably, highly effective clinical agents such as 5-fluorouracil and cyclophosphamide appeared broadly inactive across both GDSC and CTRP datasets (Fig 3E,S5A-B).^2,3^ These findings demonstrate that raw AUC-based efficacy metrics do not capture real therapeutic performance across drugs.

We performed predefined sensitivity analyses varying datasets (CTRP, GDSC, PRISM),^2-4^ in vitro efficacy metrics (EC50, AUC), and clinical efficacy and exposure definitions (ORR, Cmax,u, AUC,u, TRmin,u).^13,26^ Results were directionally consistent, and the strength of association increased systematically as exposure metrics approached clinically relevant therapeutic concentrations (Fig S5-6).

### In vitro efficacy metrics strongly correlate with clinical exposure

#### Digitization of Pharmacodynamic Database

To understand why drug identity dominated the in vitro AUC patterns,^6^ we next compared AUC values with a broad set of drug-specific clinical parameters (Fig 4). We digitized four key resources widely used in pharmacology and oncology: chemical and pharmacokinetic properties (molecular weight, half-life, protein binding), pharmacodynamic parameters (therapeutic ranges), standard cancer-drug regimens (route, dose, interval), and clinically measured exposure levels under approved dosing (unbound Cmax and AUC). ^13,22,24-26^

**Figure 4.**
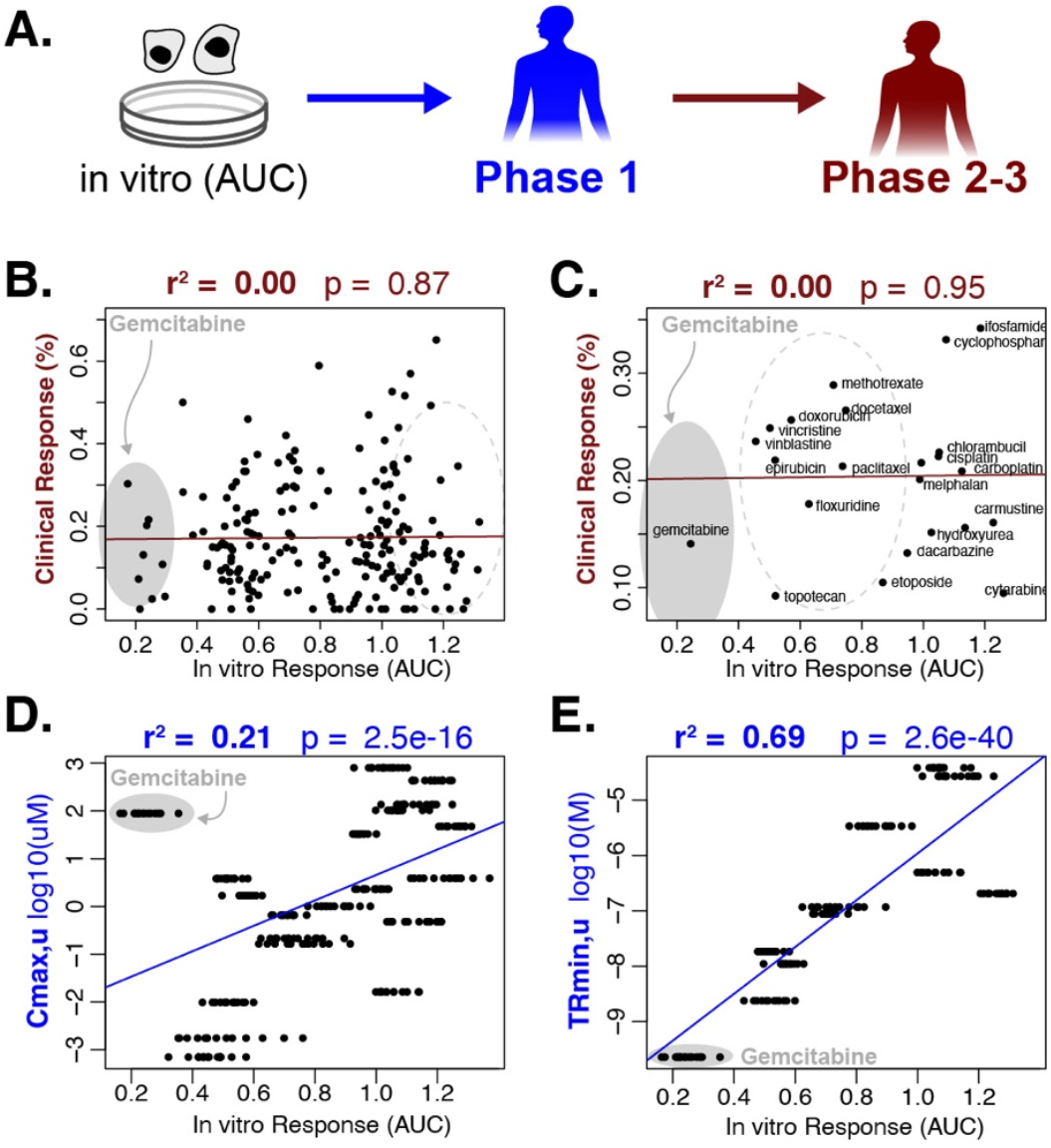
Drug-specific biases in laboratory potency metrics and their resolution by clinical exposure. **(A)** Translational framework: *in vitro* dose-response (AUC/IC50) informs discovery and Phase 1 PK (Cmax,u), while Phase 2-3 trials define efficacy by tumor regression (ORR). **(B)** Raw *in vitro* AUC values show no correlation with clinical response rates (ORR). **(C)** Averaged across cancer types, drug classes separate in to three groups: natural products align with clinical efficacy, antimetabolites/electrophiles are underestimated, and gemcitabine is overestimated. **(D)** Clinical maximum plasma concentrations (Cmax,u) partially explain drug-specific biases.^13^ **(E)** Incorporating therapeutic minimum concentrations (TRmin,u) fully restores clinical-laboratory metric concordance.^26^

Our joint pharmacokinetic-pharmacodynamic (PK-PD) dataset spans 1,352 drugs across 18 parameters and was generated by reconciling three published digital resources including a 2012 compilation of therapeutic and toxic concentration ranges across roughly 1,000 drugs and xenobiotics (Supp Data #4).^24-26^ Our clinical-exposure dataset was digitized from a 2017 print compilation of clinically relevant concentrations for 142 cancer drugs (Supp Data #5).^13^ Our cancer-regimen dataset was derived from a 2021 Cancer Therapeutics Handbook (Supp Data #6).^22^

### AUC reflects *exposure* not efficacy

Examining raw AUC and ORR across all 540 drug-cancer pairs allowed us to quantify how in vitro, Phase 1, and Phase 2-3 metrics relate (Fig. 4). Raw AUC showed no correlation with ORR (Fig. 4B; R2 = 0.00 [0.00-0.01]). Drugs separated into three groups (Fig. 3C): natural products with intermediate AUCs but strong clinical activity, antimetabolites and electrophiles that appeared inactive in vitro despite broad clinical utility, and gemcitabine as a striking outlier with very low AUCs yet only modest ORRs. These drug-specific patterns explain why raw AUC, even averaged across cancer types, cannot predict therapeutic benefit.

We hypothesized that these distortions reflect *clinically achievable exposure* not efficacy. Many antimetabolites and electrophiles reach unbound plasma concentrations of 100-400 µM in patients, far above the 5 µM ceiling of CTRP, GDSC, and PRISM assays. For example, unbound Cmax (Cmax,u) values for 5-fluorouracil, ifosfamide, and carboplatin are ∼426 µM, ∼431 µM, and ∼135 µM.^13^ Their apparent inactivity in vitro is therefore an assay-range artifact rather than true inefficacy. Consistent with this, AUC correlated significantly with clinical Cmax,u and AUCu (Fig. 3D; R^2^: 0.21 [0.21-0.34]).^13^ Gemcitabine remained anomalous, featuring both very low AUCs and among the highest Cmax,u values. Because Cmax,u captures only the upper therapeutic boundary, we next compared AUC with unbound therapeutic minimum concentrations (TRmin,u),^26^ which showed near-perfect concordance (Fig. 3E; R^2^: 0.69 [0.64-73]), indicating that AUC primarily reflects exposure requirements rather than clinical response (Fig S6).

Together, these analyses show that in vitro dose-response metrics such as AUC track exposure limits (Cmax,u, AUCu and TRmin) rather than clinical efficacy (ORR). This distinction is critical because AUC and IC50 are often interpreted as surrogates for therapeutic effect^3,32-34^ even though they mainly report the concentration range required for drug activity.

#### Limitations of exposure normalization

Normalizing AUC by Cmax,u or TRmin,u restores the expected negative association with clinical response, but correlations remain modest (Fig S7). This is likely due to the fact that concentration-centric in vitro metrics fundamentally differ from time-centric clinical endpoints (Fig 1A-B). This is evident in Fig. S7: most normalized drug-cancer pairs cluster in the lower-right quadrant, reflecting a *high false-positive rate* in which many compounds appear active *in vitro* yet have limited clinical benefit. Exposure normalization helps reduce false negatives for agents such as ifosfamide and cyclophosphamide, whose clinical concentrations far exceed standard assay ranges,^13^ but it does not correct false positives such as gemcitabine, which retains disproportionately strong *in vitro* activity despite modest clinical efficacy.^26^

High false-positive rates are a well-known property of *in vitro* drug-response systems more broadly. Two-dimensional cell cultures are inherently more drug-sensitive than tumors because they lack stromal support, have higher proliferation rates, and experience uniform drug penetration.^35-38^ Patient-derived organoids can reduce FP-rates by 10-40%,^39-43^ however they also exhibit similar limitations: classical meta-analyses and recent organoid studies report false-positive rates of roughly fifty percent (vs false-negative rates around ten percent).^44,45^ Together, these findings show that while CRR-style exposure normalization improves alignment, it cannot fully bridge the gap between simplified concentration-response assays and real clinical outcomes.

### Comprehensive Biomarker Evaluation

#### Need for Chemotherapy Biomarkers

The thirty cytotoxic chemotherapies analyzed in this study continue to form the backbone of metastatic cancer treatment today (typically combined with targeted and immunotherapies).^30,46^ Current practice applies these agents using a population-level approach, a strategy increasingly recognized as insufficient.^47^ Several groups have therefore proposed that personalizing cytotoxic chemotherapy could represent a major and cost-effective paradigm shift in oncology.^30,46,48^ Although hundreds of candidate biomarkers for chemotherapy sensitivity have been identified in laboratory studies, none have entered routine clinical use.^29^ In non-small-cell lung cancer, for example, only thirteen of eighty reported chemotherapy biomarkers have been tested clinically and only four show any diagnostic promise.^30^ However, the lack of clinical evaluation has made it difficult to compare, prioritize and combine these biomarkers.^9,30^

#### Curation and analysis of literature-derived biomarkers

Based on our framework, biomarker comparisons within a single drug are expected to be less affected by the laboratory-clinical divide than multi-drug prioritization studies. A secondary goal of this study was therefore to evaluate both clinical and in vitro correlations for all known literature-derived biomarkers across our thirty-drug library. We compiled 234 biomarkers from DrugBank and expanded this set through targeted literature searches (see Methods).^27,28^ These biomarkers fell into two major classes: sensitivity markers and resistance markers. These could be further subdivided into ***direct and indirect drug targets*** and ***drug-metabolism or transport genes***, respectively (Fig. 5A, Fig. S8). Correlations were quantified using univariate linear models comparing biomarker mRNA expression^3,32^ with clinical or laboratory drug-response metrics, as illustrated in Figure S9-10 and detailed in the Methods.

**Figure 5.**
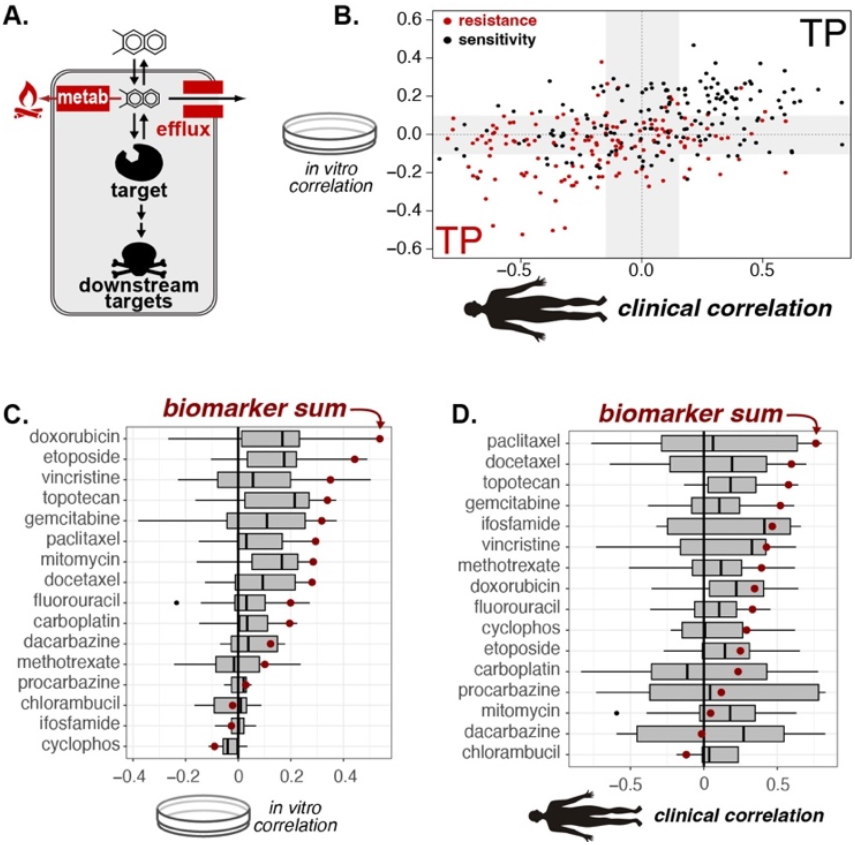
Laboratory biomarkers sets effectively predict *in vitro* and clinical drug-response. **(A)** Conceptual summary of all cytotoxic chemotherapy biomarker classes within DrugBank **(B)** Scatter plot of correlations of biomarker expression (RNAseq) with drug-sensitivity across clinical and laboratory datasets. Each point represents a drug-biomarker pair. Quadrants were defined to visualize odds ratios of literature and clinical classification agreement **(C)** Drug-specific Literature/Laboratory agreement scores across all biomarkers. **(D)** Drug-specific Literature/Clinical agreement scores across all biomarkers. Correlation of Biomarker sums are marked with a red point

#### Evaluation of biomarker-response correlations

We next evaluated biomarker correlations with drug response in vitro and in vivo across all 314 biomarker-drug pairs, plotted along the x- and y-axes of Figure 5B. Clinical and laboratory correlation directions were generally consistent (R^2^ = 0.17 [0.10-0.25], p = 2.8e-14), with most resistance markers falling in the double-negative quadrant and most sensitivity markers in the double-positive quadrant, reflecting true-positive classification for their respective categories. To quantify classification fidelity, we performed Fisher’s exact tests comparing literature-based sensitivity and resistance labels with the sign of each correlation. These classifications were significantly conserved in both laboratory data (odds ratio 3.7, p = 2.36e-8) and clinical data (odds ratio 3.9, p = 6.52e-9). Restricting analyses to stronger correlations greater than 0.1 further improved performance, yielding odds ratios of 13.3 and 8.4, respectively (Fig 5B white quadrants). Overall, sensitivity and resistance designations were broadly conserved across laboratory and clinical correlations between mRNA expression and drug response for all 314 biomarker-drug pairs.

#### Drug-specific biomarker performance

To better understand which drug classes supported the strongest biomarker performance, we next visualized biomarker-response correlations for each drug using the classification agreement score shown in Figure 5C-D. This score multiples -1 to resistance-marker correlations, so that positive values indicate agreement between the expected literature sign and the observed correlation. Using this metric, we found that most biomarkers for the top drugs showed consistent agreement between laboratory and clinical correlations. Anthracyclines, taxanes, and vinca alkaloids, in particular, demonstrated some of the strongest biomarker concordance (Fig S10).

Across drug classes, resistance markers showed stronger and more stable correlation patterns than direct drug targets (Fig S10). Multivariate models further improved agreement with literature classifications, confirming that multiple biomarkers contribute independently to chemotherapy response (Fig S11). These findings are consistent with prior modeling studies showing that simple linear models can perform well for predicting chemotherapy sensitivity.^3,32,33^

#### Performance of biomarker sums

More broadly, we found that simple sums of absolute mRNA expression values ([***E***]_*t*_) across N-biomarkers per drug produced robust predictors across most drugs, and were especially effective for the anthracyclines, taxanes, and vinca alkaloids (red dots in figure 5C-D):

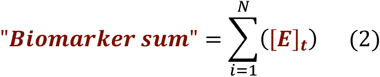

This performance can be rationalized mechanistically: under pseudo-first order Michaelis-Menten kinetics,^49-51^ the expected contribution of metabolism and transport processes to chemotherapy resistance becomes a weighted sum of pathway components, as derived in the Supporting Information:

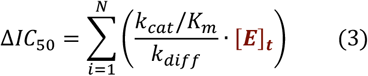

Together, these analyses show that most biomarkers maintain their expected literature-defined classification in both laboratory and clinical settings, and that biomarker sums provide unexpectedly strong performance across many cytotoxic chemotherapies, particularly those whose response is strongly shaped by drug metabolism and transport (Fig S10-11). This result is consistent with an emerging consensus that, for cytotoxic chemotherapies: no single biomarker is adequate and that multi-marker signatures will likely be required to generate reliable predictors of chemotherapy response.^10,30,52^

## DISCUSSION

The NCI1970-Meta dataset assembled here provides the first comprehensive, digitized map of single-agent clinical activity for the major cytotoxic chemotherapies. By reconciling nearly five decades of scattered Phase II-III monotherapy trials, we created a clinical anchor that enables rigorous, quantitative comparison between laboratory drug-response assays,^2-4^ pharmacokinetic exposure boundaries,^13,22,24-26^ and biomarker-response relationships.^27,28^ These analyses reveal that many long-standing translational inconsistencies can be resolved once clinical exposure is considered explicitly, and they highlight practical steps for improving preclinical drug screening and biomarker development.

### Reframing the laboratory-clinical divide

A central finding is that laboratory and clinical datasets capture *different dimensions* of drug action. Clinical response rates are dominated by cancer lineage, reflecting intrinsic differences in chemosensitivity among hematologic, epithelial, and mesenchymal malignancies. In contrast, raw in vitro potency metrics, especially AUC and IC50, are dominated by drug identity rather than cancer type. This divergence arises because commonly used in vitro assays operate within restricted concentration windows^2-4^ that exclude the true therapeutic ranges of many agents.^13^ Drugs such as 5-fluorouracil, ifosfamide, and carboplatin reach unbound plasma levels orders of magnitude higher than typical assay ceilings.^13^ As a result, they appear broadly inactive in vitro despite clear clinical efficacy.

Our analysis demonstrates that this distortion is best understood as an *exposure problem*, not a biological inconsistency. Raw AUC correlates strongly with clinically achievable concentrations (Cmax,u and TRmin,u), indicating that in vitro metrics primarily report the concentration required for inhibition, rather than the likelihood of clinical response (Fig 4, S5-6). Incorporating exposure into potency normalization restores the expected alignment between in vitro measurements and clinical outcomes (Fig S7), resolves drug-specific anomalies such as gemcitabine, and provides a coherent framework that spans discovery assays, Phase I PK studies, and Phase II-III efficacy trials.

### Implications for drug-screening and drug-repurposing

These results have immediate methodological implications. *First*, limiting assays to 5-10 µM generates systematic false-negatives for agents whose exposure in patients extends well beyond this range. Expanding screens to the full clinically relevant concentration window (0-Cmax,u) would reduce these errors and greatly improve the fidelity of training data used in machine-learning and large-scale prediction efforts.^3,32,33^ Second, comparisons across drugs require potency metrics that reflect clinical feasibility. Simple exposure-normalized measures (e.g. IC50/Cmax,u) avoid over-prioritizing compounds with low achievable plasma levels and provide more realistic ranking of candidate agents in multi-drug screens. This approach is already recommended by the FDA for evaluating drug-drug interactions,^12^ and increasingly used in patient-derived organoid drug-testing.^15-18^

### Clarifying biomarker interpretation

A second major contribution of this dataset is a systematic, side-by-side comparison of 234 literature-derived biomarkers across both in vitro and clinical endpoints. By evaluating the sign and magnitude of biomarker-response correlations for 314 drug-biomarker pairs, we show that most sensitivity and resistance labels are conserved across laboratory and clinical contexts, particularly for anthracyclines, taxanes, and vinca alkaloids (Fig 5). Drug metabolism and transport pathways performed especially well, consistent with their direct influence on intracellular drug levels (Fig S9-11).

Simple sums of biomarker expression produced unexpectedly strong performance across many drugs (Equation 2). This is mechanistically intuitive: when drug inactivation, efflux, and target abundance act under pseudo-first-order kinetics, their combined effect approximates a weighted sum of contributing components (Equation 3). These findings support growing evidence that multi-gene signatures outperform single biomarkers and that chemotherapy response is best understood as the aggregate of multiple, modest-effect pathways.^10,30,52^

### Limitations of the study

The clinical trials underlying NCI1970-Meta were conducted between 1955 and 2000 and reflect the patient populations, diagnostic criteria, and trial practices of that era (Fig 2). Nevertheless, they remain the only large-scale record of cytotoxic monotherapy efficacy, as modern trials overwhelmingly evaluate combinations. Pharmacokinetic benchmarks were derived from therapeutic drug-monitoring studies and represent population averages; variability between patients, regimens, and formulations is an inherent source of uncertainty.^13,22,24-26^ Finally, because clinical efficacy depends on tumor evolution, immune interactions, microenvironmental constraints, and host toxicities, exposure normalization should be viewed as a necessary but not sufficient step toward clinical prediction. Future work will need to integrate integrating that incorporate tissue-exposure, tumor state, and microenvironment in a unified mathematical form.

## CONCLUSION

By digitizing the historical clinical evidence for 30 major chemotherapies and aligning it with high-throughput in vitro datasets, clinical exposure benchmarks, and a comprehensive compendium of biomarkers, we provide a resource that directly addresses one of the most persistent gaps in translational oncology. These analyses show that preclinical drug-response assays do contain the biological signals seen in patients: but these signals are interpretable only when grounded in clinical exposure. In this framework, both laboratory pharmacology and biomarker studies gain clarity, enabling more accurate drug prioritization, more biologically informed biomarker discovery, and a more principled connection between laboratory potency and patient benefit.

## ACKNOWLEDGEMENTS

E.F.D. acknowledges Matthew Clarke for helpful discussion and the University of Georgia for financial support.

## AUTHOR CONTRIBUTIONS

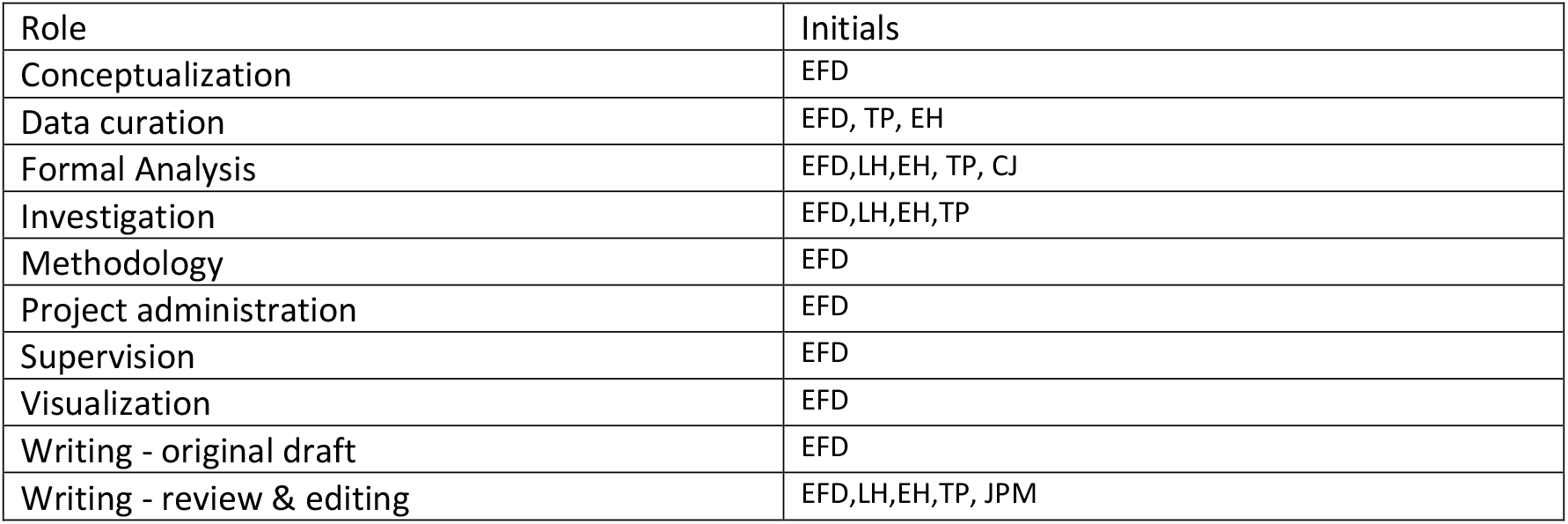

## METHODS

### Computational Method Details

#### CURATION OF SINGLE-AGENT RESPONSE RATE DATA (Figure S1-4 and Figure 2)

Our single-agent dataset (Figure 1A) compiles first-line metastatic cancer response rates from phase II and III trials published between 1950 and 2000. **Data sources and scope:** The dataset was assembled from the *NCI Textbook: Single Agents in Cancer Chemotherapy*^*21*^ and 48 meta-reviews published between 1970 and 2000 (Figure S1). Fully curated data are provided in Supplementary Data 01, with each curation step detailed in Supplementary Tables 02-03 and graphically summarized in Figure S02.

**Eligibility criteria** were defined using the inclusion rules of the NCI monograph, requiring first-line treatment of metastatic disease, single-agent phase II or III design, and tumor response evaluation using WHO-style definitions. Although WHO criteria were formally codified in 1978,^53^ the internal NCI definitions applied in earlier decades used equivalent thresholds for the distinction between partial response and stable disease. Because complete response (CR) was not consistently defined across decades, CR rates were not extracted or used in any analysis. To enable meaningful comparison with modern datasets, additional criteria were introduced, excluding early-era agents that are not considered oncology therapeutics today and restricting included drugs to those listed in the *Physicians’ 2021 Cancer Chemotherapy Manual*.^22^ Analyses were limited to first-line treatment settings to maintain consistency across eras.

The structure of the underlying monograph introduces additional constraints on the dataset. The NCI cooperative trials program, which began in 1955, generated a coordinated national portfolio of phase II single-agent trials, shaping both the density and timing of available data. The monograph systematically excluded combination chemotherapy, adjuvant or regional perfusion studies, and most foreign-language reports, thereby restricting the collected evidence to systemic, intravenous or oral single-agent therapy. Evaluability criteria were not standardized across investigators (i.e. ‘patients evaluated’ frequently differed from ‘patients treated’) and the definition of ‘objective response’ varied, though in solid tumors the compilers generally required >50% measurable tumor regression. Definitions of complete response, particularly in leukemias, differed widely (e.g., M1 marrow vs. full hematologic recovery), motivating our decision not to extract or analyze CR rates. Tumor inclusion was asymmetric: breast, colorectal, lung, melanoma, lymphomas, and leukemias were consistently reviewed, whereas other cancers were included only when adequate data existed (Figure S4. These inherited structural features, along with substantial dose-schedule heterogeneity across studies and cooperative groups, contribute patterned variance that is addressed in downstream analyses.

#### Epoch definitions and rational

All included trials predated RECIST, and their response assessments fell entirely within the pre-WHO or WHO frameworks. To contextualize temporal heterogeneity, trials were grouped into three predefined epochs: the pre-WHO era (1955-1978), in which most classical cytotoxics were developed; the WHO era (1979-1988), dominated by platinum agents and camptothecins; and the late WHO/pre-RECIST era (1990-2000), characterized by taxanes and gemcitabine (Figure 2). Most single-agent drug development occurred entirely within a single epoch, enabling internally consistent biomarker- and cancer-level analyses. In contrast, all cross-drug comparisons necessarily spanned multiple epochs, introducing additional variance that is addressed in downstream statistical modeling.

#### Data extraction and curation

Raw response numbers from the NCI monograph were manually encoded into a four-column matrix (Figure S1A) in Excel (Supplementary Data 02: “NCI1970-update > NCI 1955-70”). Matrix conversion revealed high sparsity (58%) (Figure S1B), which limited early statistical analyses. To reduce missingness, we consolidated lymphoma, leukemia, and sarcoma data and integrated response data from 48 additional meta-reviews published between 1970 and 2000 (Figure S1C). These additions substantially improved dataset completeness and enabled the cross-cancer, within-drug, and between-drug analyses described in the main text.

#### Meta-review integration and data finalization

The meta-reviews included in this study were assembled as direct successors to the 1970 NCI monograph. Twelve were authored by the original monograph investigators (Stephen K. Carter and Robert B. Livingston), and thirty-six additional reviews were identified through citation chains stemming from either the monograph itself or its subsequent follow-up publications.

These reviews generally aimed to update activity estimates for newly introduced drugs over the next three decades or to expand coverage for cancer types under-reported in the original volume. Although broadly complementary, the meta-reviews contained overlapping content with NCI1970 and with each other. This overlap required two complementary curation strategies because many meta-reviews reported only final aggregated response rates without full trial-level detail.

In the first strategy, we recovered all available primary-source citations, explicitly removed duplicates based on exact drug-cancer-reference matches, and constructed a deduplicated primary-source dataset suitable for random-effects meta-analysis (Supplementary Table 03). In the second strategy, we encoded the aggregated response rates from every meta-review and applied a rule-based selection algorithm to identify, for each drug-cancer pair, the largest evaluable cohort ever reported across all reviews. Because these later aggregates often lacked full trial-level provenance, they could not be meta-analyzed, but they provided the most statistically powered consensus estimates available (Supplementary Table 02).

Meta-reviews were encoded in a uniform tabular format organized by drug (NCI1970-update > drug-meta) or cancer type (NCI1970-update > NCI cancer-meta), and inclusion criteria matched those of the monograph: single-agent phase II data, WHO-style response definitions, and first-line treatment of metastatic disease. Our hierarchical update algorithm (Figure S2B) replaced NCI1970 entries whenever a larger and higher-powered dataset was available, filling many gaps and adding 25 additional drugs. The resulting consolidated dataset, NCI1970-Meta, reduced sparsity from 58% to 41%. Final response estimates were cross-referenced against modern oncology textbooks to ensure concordance with established clinical summaries(e.g. pages 548, 637, 748, 783, 823, 858, 1035, 1065, 1192, 1305, 1431, 1550 in the 11^th^ edition of *Cancer - Principles & Practice of Oncology*).^29^

#### Random-effects meta-analyses

For drug-cancer pairs in which primary-source trials were recoverable (Supplementary Table 03), we conducted independent random-effects meta-analyses using all non-duplicate studies reporting single-agent response rates. For each trial, we extracted the number of responders and total evaluable patients, and analyzed each drug-cancer pair using a binomial-normal model with logit-transformed proportions and a REML estimator, allowing us to account for both within-study sampling variance and between-study heterogeneity (τ2). Zero-event studies were retained without continuity correction using the PLO transformation implemented in the *metafor* package. For every meta-analysis, we report the pooled ORR, 95% confidence interval, number of contributing studies, τ2, and I2. All pair-level results are provided in full in Supplementary Table 03: including: including trial counts, pooled estimates, and heterogeneity metrics.

### CURATION OF FDA-INDICATION DATA (Figure 3C)

To capture contemporary “standard of practice” information for the 34 drugs and 19 cancer lineages represented in our dataset, we manually encoded FDA-approved clinical indications using the 2021 Physicians’ Cancer Chemotherapy Drug Manual.^22^ For each drug-cancer pair we constructed a blank matrix and initialized all entries to “0,” then replaced entries with “1” when a drug held an FDA indication for that cancer type (Supplementary Data 01: Figure Data > FDA). Because the FDA manual lists indications at the level of common clinical entities, we first reconciled each indication to a unified lineage taxonomy based on organ site and histology, and then mapped each lineage to its corresponding TCGA cancer-type abbreviation.^54^ For solid tumors, this lineage harmonization was straightforward; however, lymphoma and leukemia categories were aggregated globally because the NCI1970 monograph did not distinguish subtypes at the granularity used in modern TCGA classifications.^*21*^

FDA indications necessarily reflect later-era clinical practice, most often representing the incorporation of these drugs into combination chemotherapy regimens developed well after the completion of early single-agent trials. In contrast, the ORRs compiled in our dataset are tied to discrete historical epochs reflecting the time periods in which single-agent evaluations for each drug were performed. We therefore matched lineage assignments consistently but treated temporal information separately: lineage reconciliation was anchored to modern TCGA categories, whereas ORR values remained epoch-specific. Because combination regimens were universally designed under the assumption that constituent agents possessed intrinsic single-agent activity,^55^ the alignment between modern FDA combination indications and historical single-agent ORRs provides a meaningful test of compatibility between early monotherapy response profiles and later real-world therapeutic use.

### IN VITRO DRUG SENSITIVITY DATASETS (Figure 3E, S5-7)

To evaluate how laboratory drug-response metrics relate to clinical activity, we curated CTRP, GDSC, and PRISM from the DepMap portal (https://depmap.org/portal/download/) release 20Q2.^56,57^ Using DepMap was essential because it provides harmonized drug names, cell-line identifiers, and lineage annotations across all screening datasets, preventing inconsistencies that commonly arise when these resources are downloaded separately. All molecular biomarker data (RNA-seq, copy number, and mutations) were obtained from CCLE (DepMap 20Q2), ensuring that pharmacologic and molecular features were aligned to the same standardized cell-line nomenclature.

AUC values were used as the primary measure of in vitro activity because they capture overall efficacy across the full dose-response range and are less sensitive to curve-fitting variability than EC50/IC50. EC50 values were analyzed in parallel as a sensitivity check; they correlated strongly with AUC across all datasets, and all study conclusions remained unchanged, confirming that results were not metric-dependent. The three screening resources were used for complementary purposes. CTRP and GDSC each include ∼900 cancer cell lines spanning hematopoietic, epithelial, and mesenchymal malignancies, making them best suited for cancer-lineage and biomarker analyses. PRISM, while limited to adherent epithelial lines, contains a far larger and more clinically enriched compound library including ∼1,400 FDA-approved drugs and nearly all widely used oncology agents. All key findings were tested across CTRP, GDSC, and PRISM, and results were consistent: lineage-level signal was strongest in CTRP/GDSC, and drug-level signal was strongest in PRISM.

### PK-PD Drug Exposure Data Curation and Analyses (Figure 4, S6-7)

To determine whether in vitro drug-response metrics reflect intrinsic drug activity or simply the clinically achievable range of exposure, we compiled a multi-source pharmacokinetic-pharmacodynamic (PK-PD) dataset and compared unnormalized and exposure-normalized in vitro metrics (Supplementary Tables 4-6). All PK-PD variables were digitized from authoritative pharmacology and oncology references and harmonized by drug name to allow direct comparison with AUC and EC50 values derived from CTRP, GDSC, and PRISM.

We first assembled a broad PK-PD panel encompassing molecular properties (molecular weight, half-life, protein-binding fraction), therapeutic and toxic concentration ranges, and clinical regimen attributes (route, schedule, and standard dosing). These parameters were extracted from three published digital resources, including a 2012 reference that reports therapeutic and toxic plasma ranges for over 1,000 drugs and xenobiotics (Supplementary Data 4).^24-26^ Clinically measured exposure levels (total Cmax, unbound Cmax, AUC, and unbound AUC) were digitized from a 2017 print compendium of clinically relevant concentrations for 142 oncology drugs (Supplementary Data 5).^13^ Standard cancer-drug regimens were obtained from the *Physicians’ 2021 Cancer Chemotherapy Manual (Supplementary Data 6)*.^22^ Together, these sources generated a reconciled PK-PD dataset spanning 1,352 drugs and 18 exposure-related parameters.

Using this harmonized dataset, we compared raw in vitro potency (EC50), in vitro efficacy (AUC), and exposure-normalized metrics in which each drug’s EC50 or AUC was divided by clinically observed Cmax, unbound Cmax, AUC, or unbound AUC. These exposure-normalized values were then correlated with clinical ORR values derived from the NCI1970-Meta dataset. This analysis allowed us to quantify whether in vitro sensitivity metrics primarily reflect intrinsic biological activity or instead track the magnitude of clinical drug exposure achievable in patients. Across drugs and cancers, raw AUC values were strongly associated with clinical exposure parameters but only weakly associated with clinical ORR after exposure normalization, indicating that in vitro AUC is heavily shaped by pharmacokinetic ceilings and only partially predictive of therapeutic activity.

### BIOMARKER-EXPRESSION DATASETS (Figure 5, S8-11)

Biomarkers associated with drug sensitivity or resistance for our 34-drug library were curated from DrugBank (RRID:SCR_002700; https://go.drugbank.com/releases/latest).^27,28^ and from >200 primary literature sources (Figure S6). This yielded 41 drug targets, 47 drug transporters, 62 drug-metabolism enzymes, and 84 additional downstream biomarkers reported to influence drug response. Biomarkers were categorized as sensitivity markers (drug targets, drug importers, or genes whose increased expression was reported to enhance drug activity) or resistance markers (drug-metabolizing enzymes, efflux transporters, or genes whose increased activity reduced drug efficacy). For downstream markers, the direction of association (sensitivity vs resistance) was assigned exactly as described in the original source. All curated drug-biomarker assignments are summarized in Figure S8.

All in vitro expression data were obtained from the CCLE (DepMap 20Q2),^56,57^ which ensured consistent gene identifiers and cell-line naming across CTRP, GDSC, and PRISM. All clinical expression data were obtained from TCGA PanCancer via cBioPortal.^54^ For both datasets, RNA-seq values were quantified as transcripts per million (TPM) and also transformed to log2(TPM + 1) for downstream correlation analyses.

Biomarker correlations with drug response were evaluated in the CTRP dataset, which provided the largest cell-line panel and the broadest coverage of drugs with defined biomarkers. Using the procedure illustrated in Figure S9, we computed correlations for 314 drug-biomarker pairs, generating the data underlying Figure 5B. Because anthracyclines, taxanes, and vinca alkaloids showed the strongest correspondence between clinical ORRs and laboratory biomarker associations, we examined these drug classes individually in Figure S10. For each class, we evaluated univariate correlations and then fit multivariate models to reconcile inconsistencies in biomarker sign, revealing that multiple biomarkers contributed independently to response.

To generate composite biomarker scores, we multiplied TPM expression values by the curated biomarker matrices to produce absolute-expression-weighted biomarker sums. TPM (rather than log-transformed expression) was used because our first-principles PK-PD model predicts that intracellular drug concentration is proportional to the absolute expression of transporters and metabolic enzymes; therefore weighting by TPM preserves biologically meaningful differences in predicted intracellular drug levels. These biomarker sums were used consistently across Figures S6, S8, and S10.

### STATISTICAL ANALYSES

All analyses were performed in R using custom scripts (https://github.com/fingolfn/nci1970). To quantify the structure of clinical and laboratory response matrices, we used two-way ANOVA without interaction terms to partition variance between drug identity and cancer lineage. This framework was applied to clinical ORR matrices and to in vitro AUC matrices from CTRP, GDSC, and PRISM, allowing direct comparison of lineage-driven versus drug-driven effects. Clinical ORR values were meta-analyzed using random-effects models, and weighted linear regression (weights = trial sample size) was used to relate ORR to FDA-indications, disease incidence, drug exposure, and in vitro potency. For all regression models, we report coefficients, p-values, R, and RMSE, and quantify uncertainty via bootstrap resampling (n = 1,000) and analytical R2 confidence intervals from the noncentral F-distribution.

To evaluate whether in vitro potency measures primarily reflect intrinsic drug activity or the limits of clinical exposure, we computed exposure-normalized metrics (e.g., AUC/Cmax, EC50/Cmax, AUC/AUCu) and analyzed them using the same regression framework. Correlations between raw and exposure-normalized metrics, as well as their associations with clinical ORR, were visualized using scatterplots with fitted lines and annotated with correlation statistics. Hierarchical clustering and heatmaps of drug-cancer matrices used standardized (z-scored) values with Spearman or Euclidean distance and average linkage to assess global structure.

Biomarker analyses were performed using univariate linear models across 314 curated drug-biomarker pairs, with sign-consistency evaluated using Fisher’s exact tests. Multivariate models were used for drug classes with strong biomarker signal (anthracyclines, taxanes, vinca alkaloids) to assess whether multiple biomarkers contributed independently to drug response. Gene-expression features were analyzed using TPM or log2(TPM+1), depending on whether absolute or relative expression was required. All key analyses— including variance partitioning, exposure-normalization, biomarker associations, and regression models—were repeated across CTRP, GDSC, and PRISM and across AUC and EC50 metrics to confirm that conclusions were robust to dataset choice and potency measure.

## Resource Availability

### Lead Contact

Further information and requests for resources should be directed to and will be fulfilled by the Lead Contact, Eugene Douglass.

### Materials Availability

This study did not generate new unique reagents.

### Data and Code Availability

Data visualization and analysis code is publicly available through a GitHub Repository (https://github.com/fingolfn/nci1970).

