## Supplementary Information for "A comprehensive pharmacodynamic dataset reconciles 50 years of laboratory and clinical knowledge"

### SUPPORTING INFORMATION:

#### Assembly of Clincial Trial Dataset

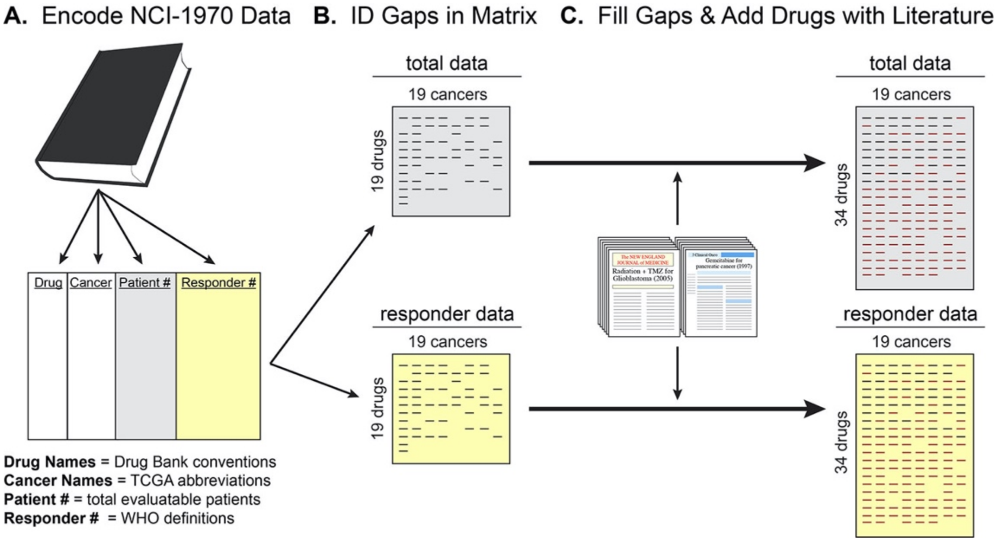

Figure S1. Encoding of Unpublished Clincial Trial Data from NCI Textbook. The clinical objective response database analyzed here was built by **(A)** encoding a 463 clinical trials published by the NCI in 1970 **(B).** Rendering the data as a matrix to enable analysis and identification of knowledge gaps. **(C)** Filling of gaps using clinical meta-review data published between 1970 and 2000.

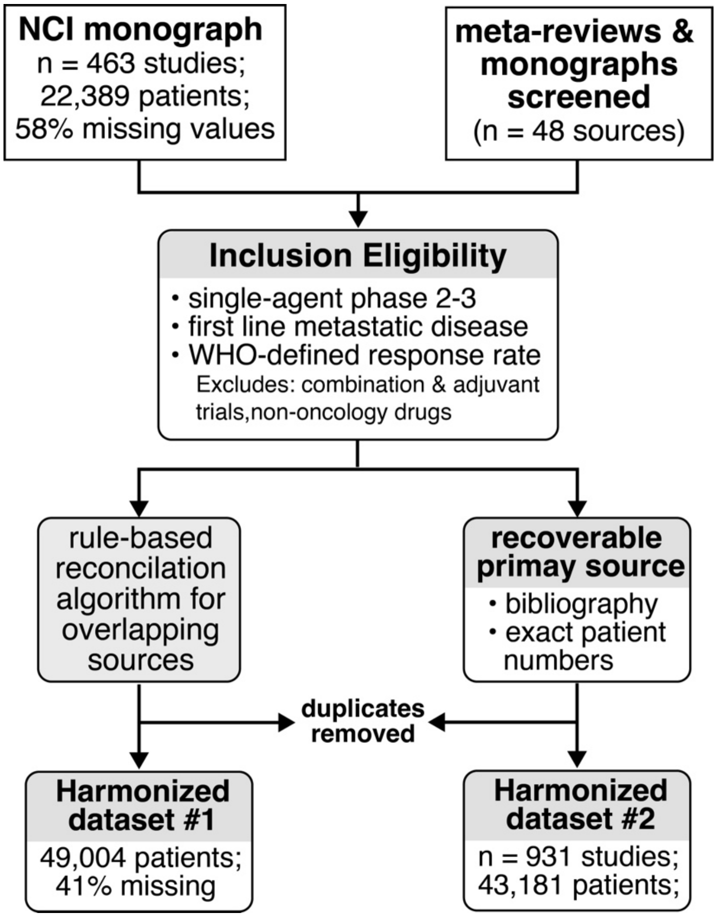

Figure S2. Flow Chart of Systematic Curation of Single-Agent Meta-Analyses papers NCI1970 textbook was integrated with data reported from 48 meta-reviews published between 1970-2000

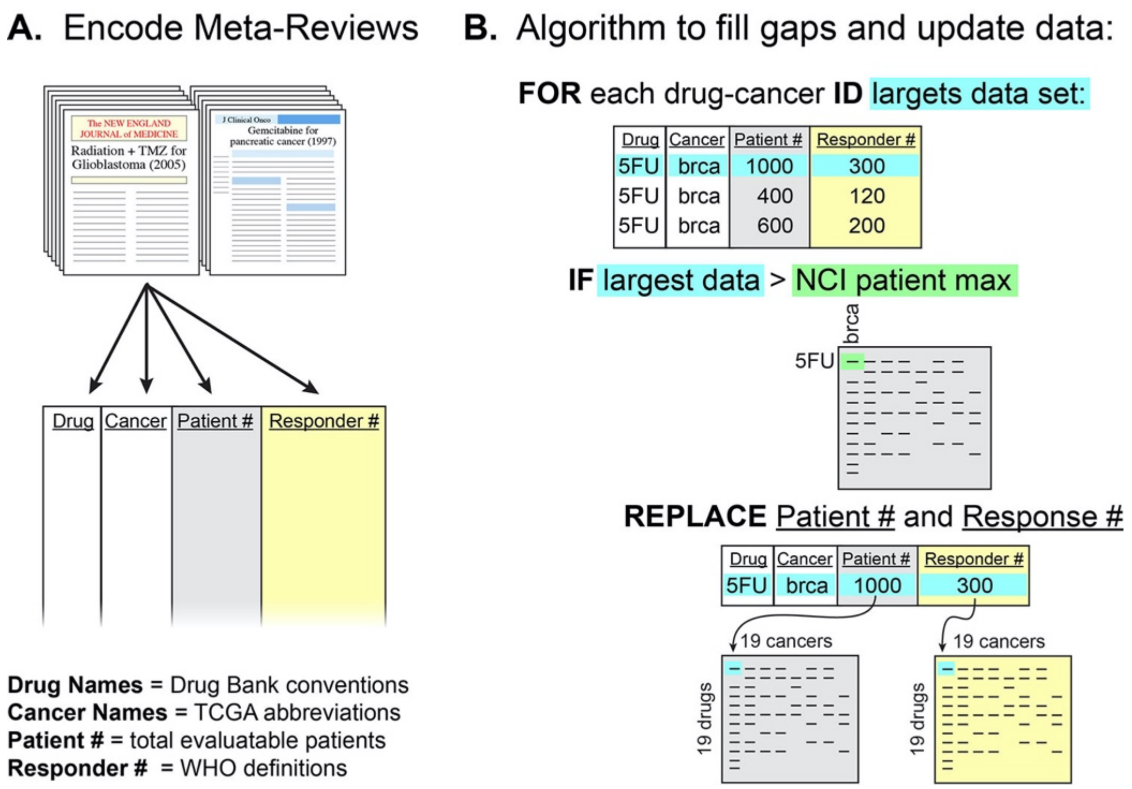

Figure S3. Algorithm to Update NCI1970 Detailed overview of updating procedure for NCI1970 with meta-reviews **(A)** First data from meta-reviews was encoded in a similar manner to raw NCI1970 data[1-48] **(B).** An algorithm was designed to: for each drug-cancer pair (1) identify the meta-review with the largest aggregate data set and (2) replace original NCI1970 matrices

#### Analyses of Bias within Clinical Data

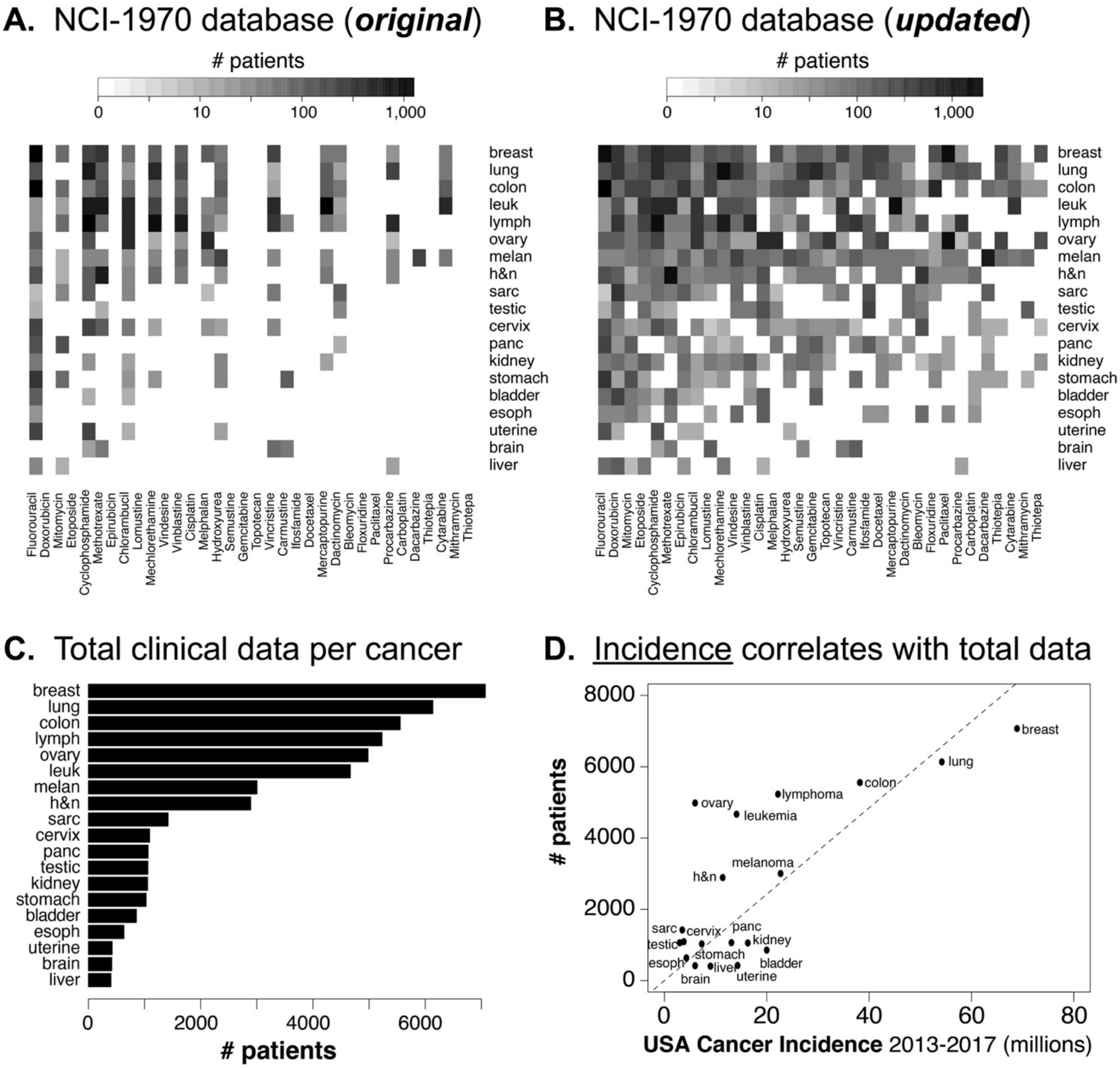

Figure S4. Overview of NCI1970-meta database. **(A).** The initial dataset was encoded directly from a textbook published in and included data for 19,958 evaluable patients across 19 cancers and 19 drugs. **(B)** This database was expanded to fill gaps and add an additional 25 drugs using metareviews published between 1970 and 2000. **(C)** Though meta-reviews for every type of cancer were identified, the number of studies varied widely by cancer-type**. (D).** Cancer incidence provides an explanation for amount of clinical data in NCI1970

#### In vitro Data Comparison with Clinical Data

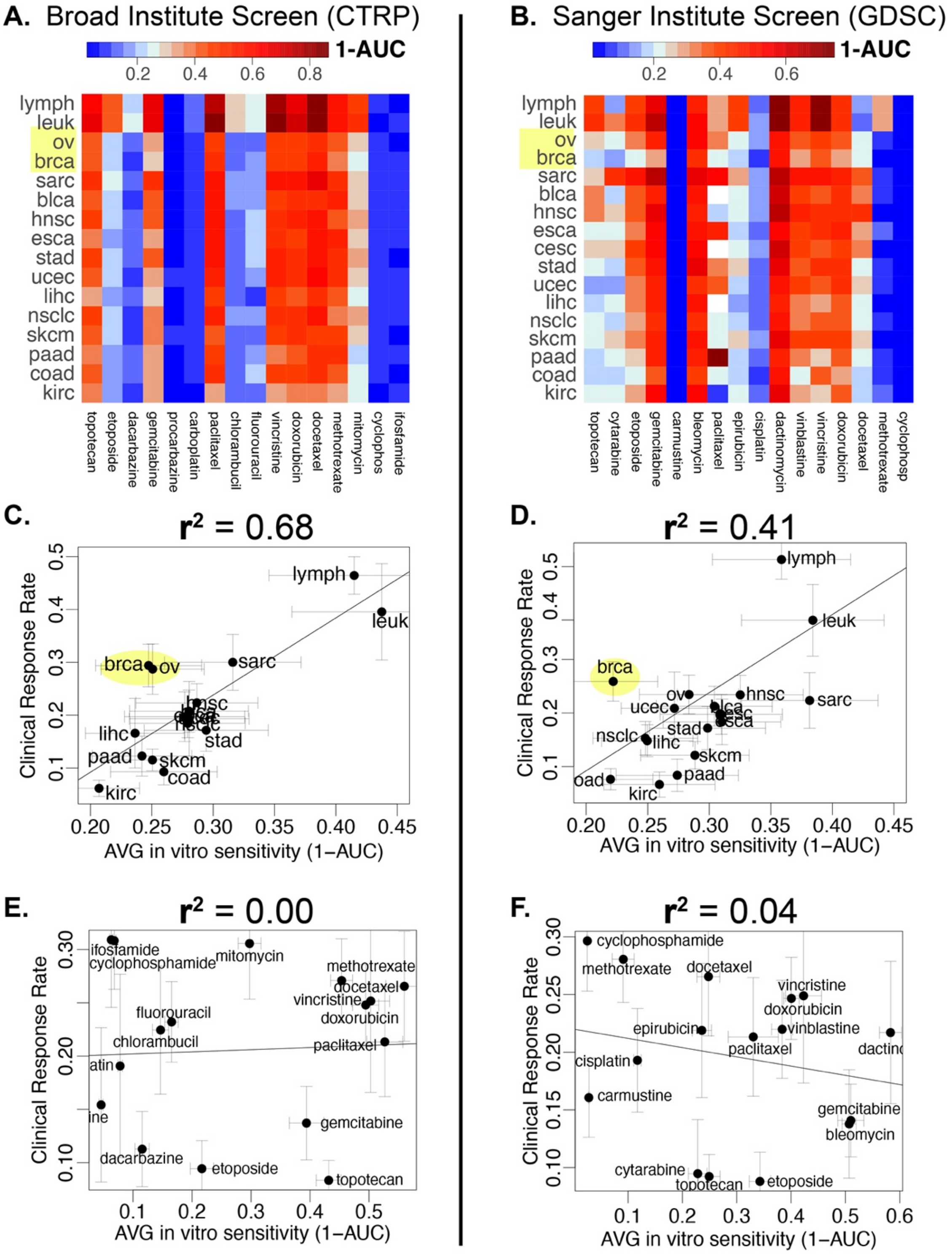

Figure S5. Comparison of Raw and Drug-Normalized Dose-response Data. (A-B) Both CTRP and GDSC in vitro drug sensitivity databases deviate from NCI1970-meta data in the very low efficacy of anti-metabolites and electrophilic chemotherapies. (C-D). Cancer-rankings of sensitivity were conserved across clinical and in vitro data (E-F) Drug-rankings showed no correlation between clinical and in vitro data

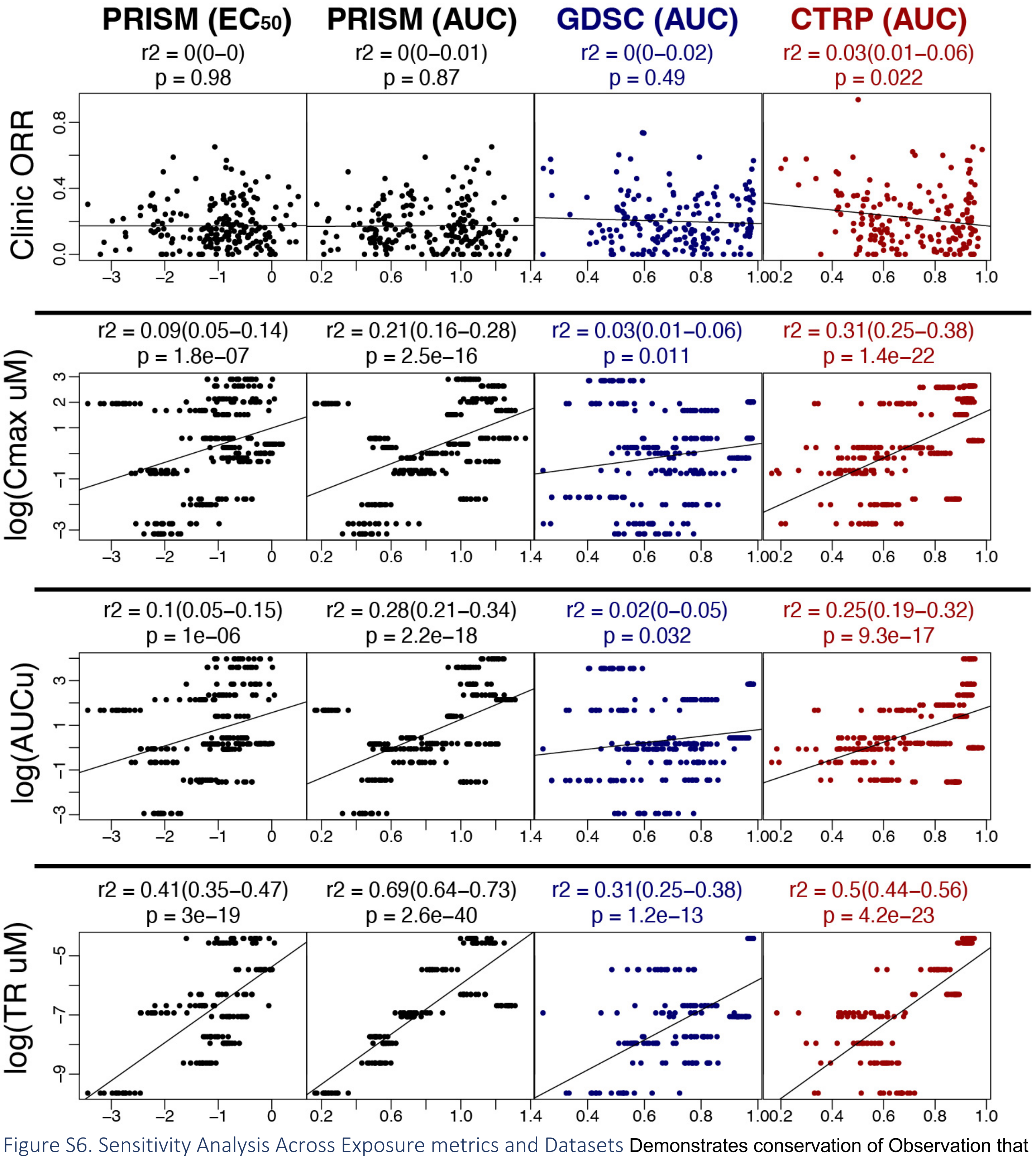

Figure S6. Sensitivity Analysis Across Exposure metrics and Datasets Demonstrates conservation of Observation that in vitro efficacy metrics correlate with therapeutic exposure not clinical efficacy

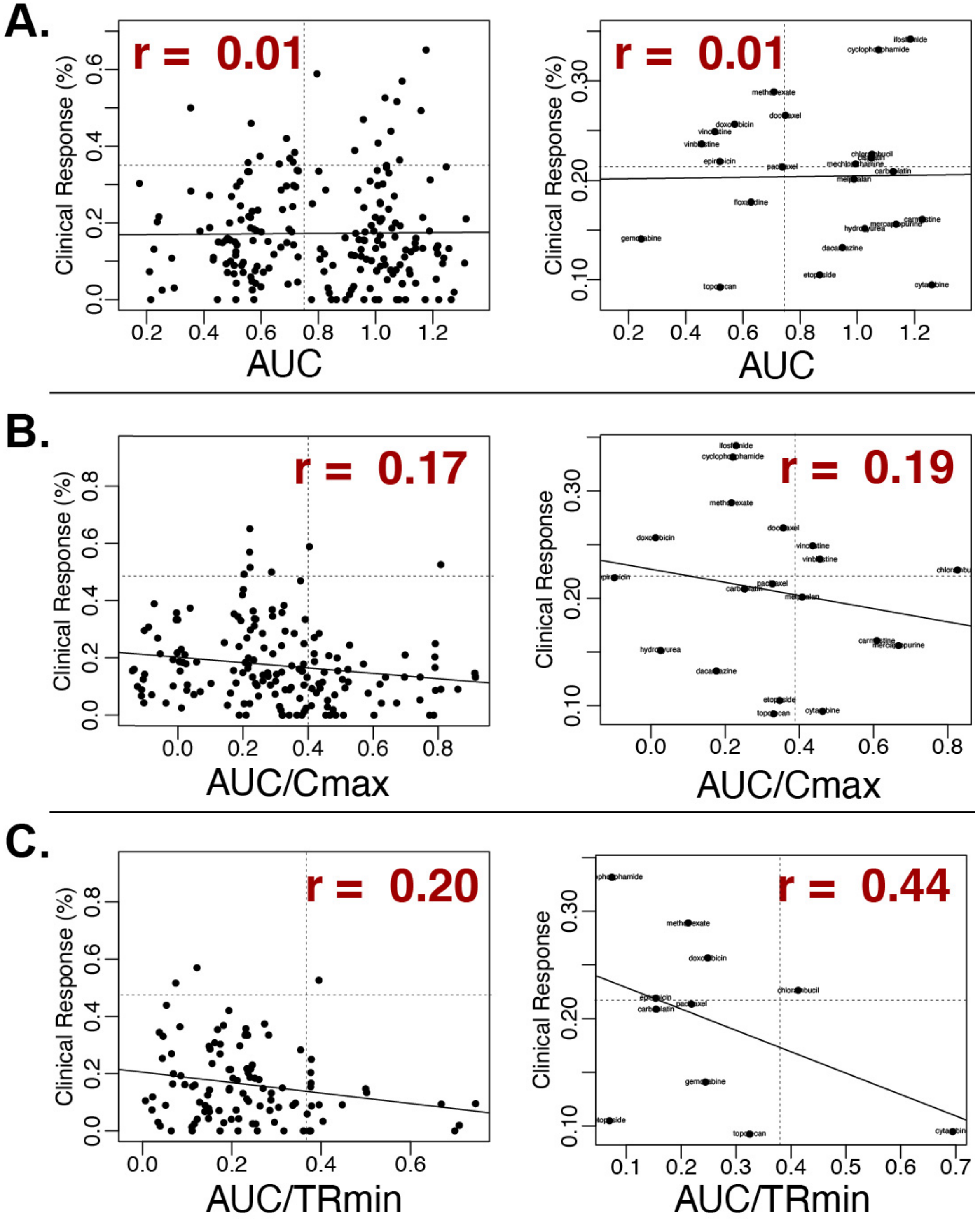

Figure S7. Drug-Normalized Dose-Response Data Correlates with Clinical Data Encouragingly in vitro data sets show significant correlation with clincial data when raw dose-response metrics (AUC) were normalized by drug-exposure (either Cmax or TRmin)

#### Assembly of Clinical Biomarker Dataset

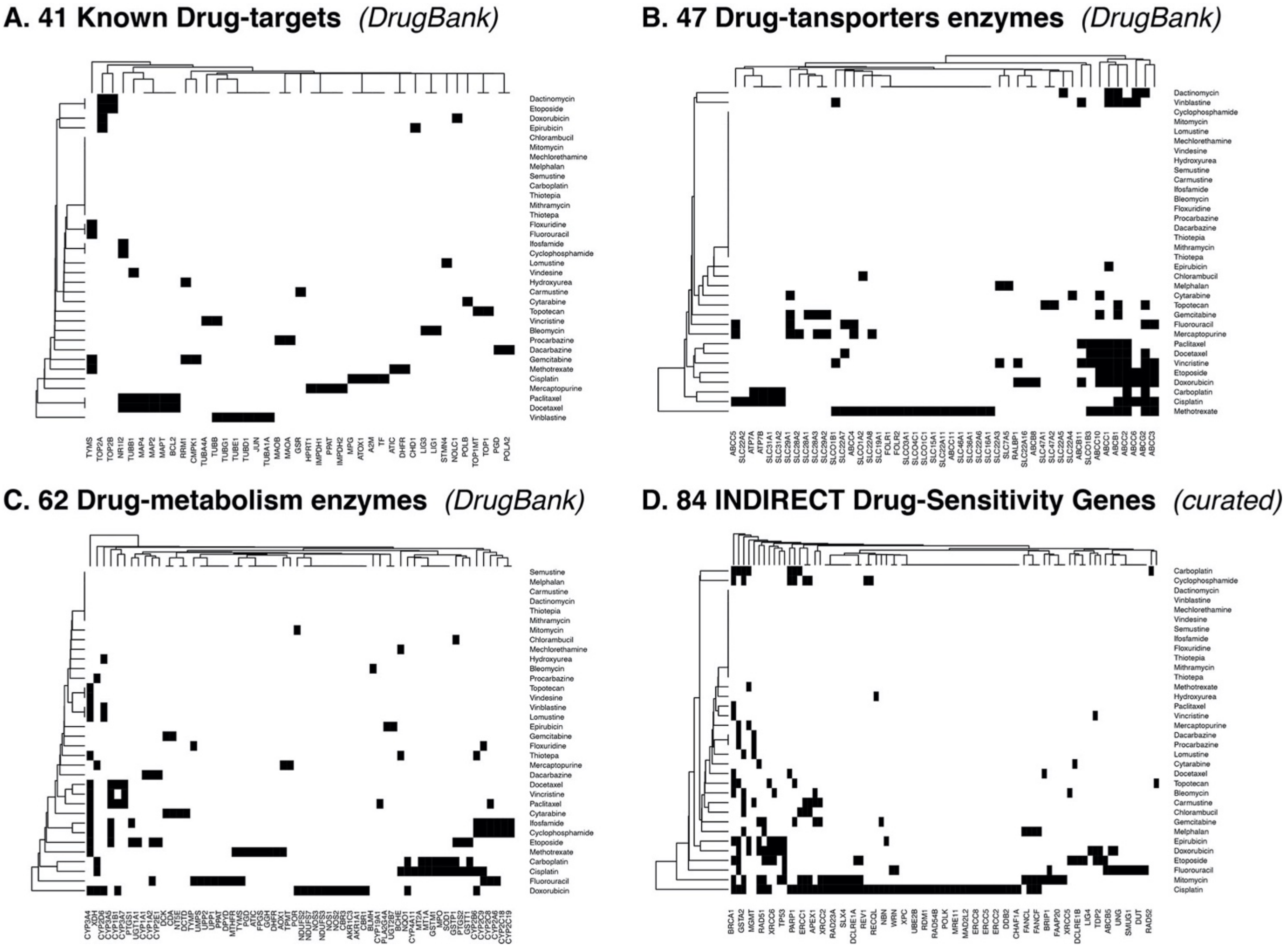

Figure S8. Biomarker Dataset Compiled from DrugBank and Manual Curation. Assembly of data-matrices documenting experimental evidence for genes that have a causal role in determining a cancers sensitivity to a particular drug including: direct drug targets **(A).** direct drug transporters **(B).** direct drug-metabolizing enzymes **(C)** and indirect pathway-based effectors of drug-efficacy **(D).** [11,12,14,21-25,27,29-37,39-41,44,45,49-256]

### Comparison of Biomarker Performance: clinical vs in vitro

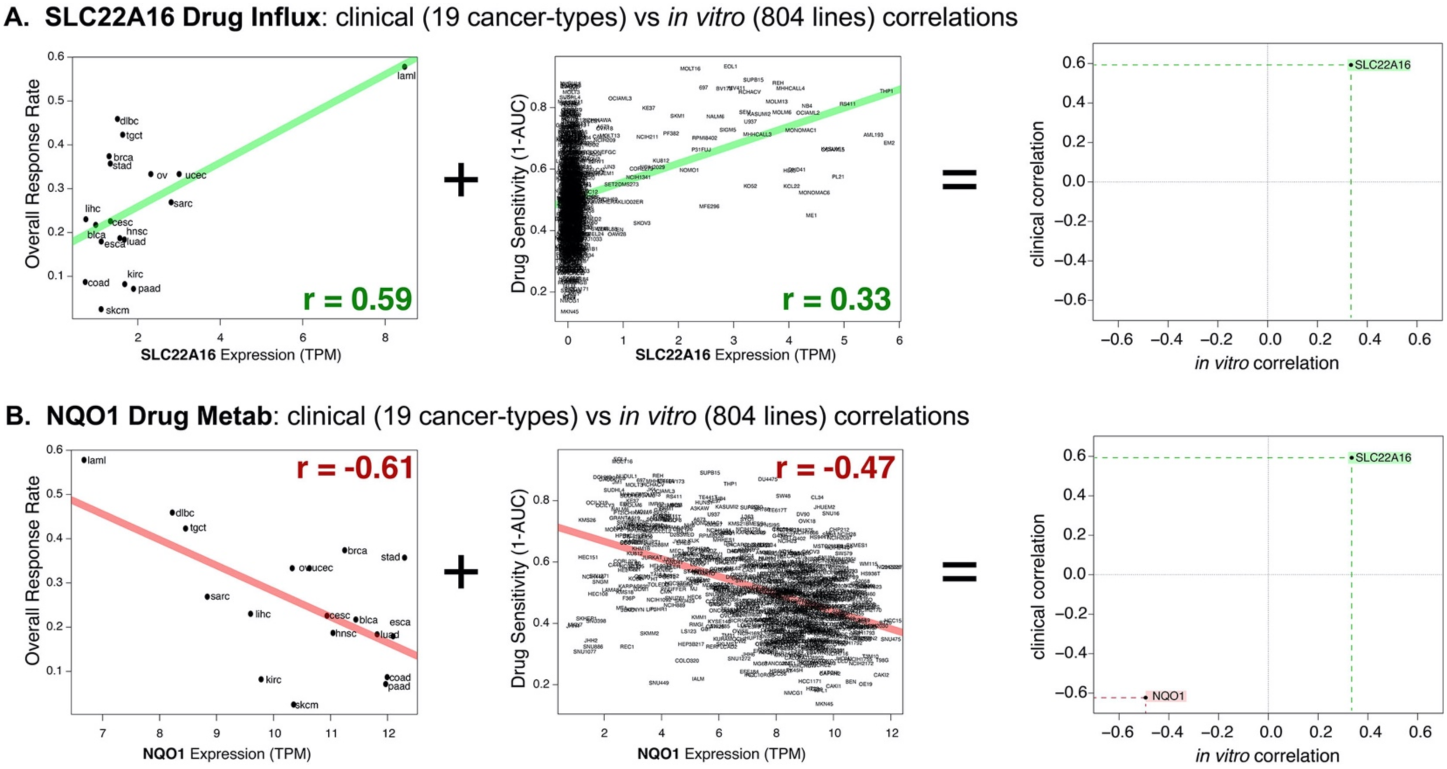

Figure S9. Example of Comparison of Individual Biomarkers Clinically and In Vitro For each experimentally validated "causal biomarker" we performed a correlation analysis on clinical data (left) and *in vitro* data right (middle) and visualized the concordance via a scatter plot **(A).** The Doxorubicin influx pump SLC22A16 shows a **positive correlation** with doxorubicin sensitivity in both clinical and *in vitro* data sets **(B).** The Doxorubicin metabolism enzyme NQO1 shows a **negative correlation** with doxorubicin sensitivity in both clinical and *in vitro* data sets

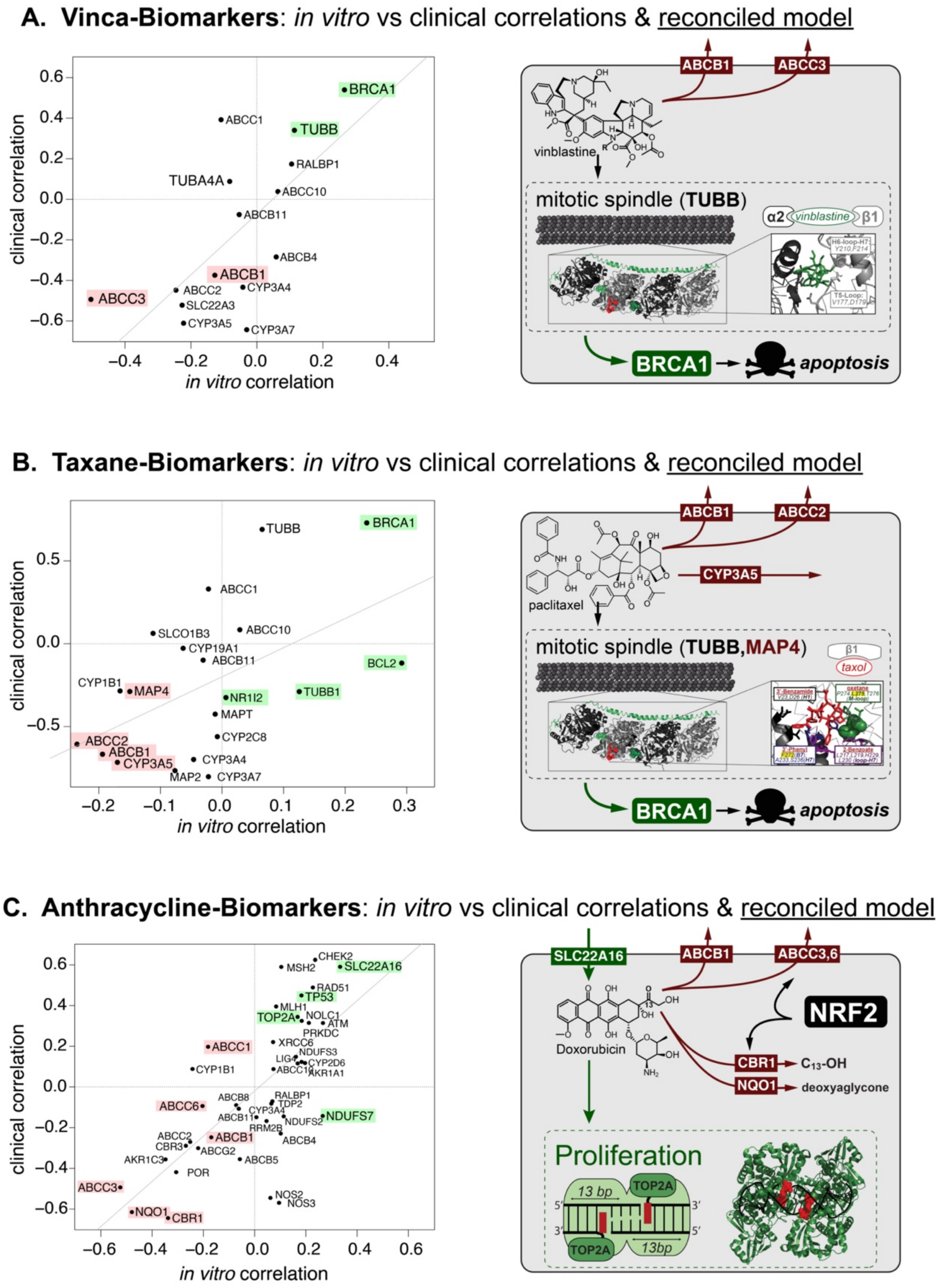

Figure S10. Overview of Correlation between Biomarkers Predictive Power. Comparison of clinical and laboratory biomarker correlations for the three most commonly indicated classes of chemotherapies. **(A).** Most biomarkers of vinca-alkaloid sensitivity show similar correlations in vitro and in the clinic. Four show independent, statistically significant predictive power when fit to a multiple-linear model. **(B)** Most biomarkers of taxane sensitivity show similar correlations in vitro and in the clinic. Eight show independent, statistically significant predictive power when fit to a multiple-linear model. Unfortunately, three of these biomarkers show inconsistent correlations in vitro and in the clinic. **(C).** Most biomarkers of anthracycline sensitivity show similar correlations in vitro and in the clinic. ten, show independent, statistically significant predictive power when fit to a multiple-linear model. Several of these are members of the antioxidant response pathway.

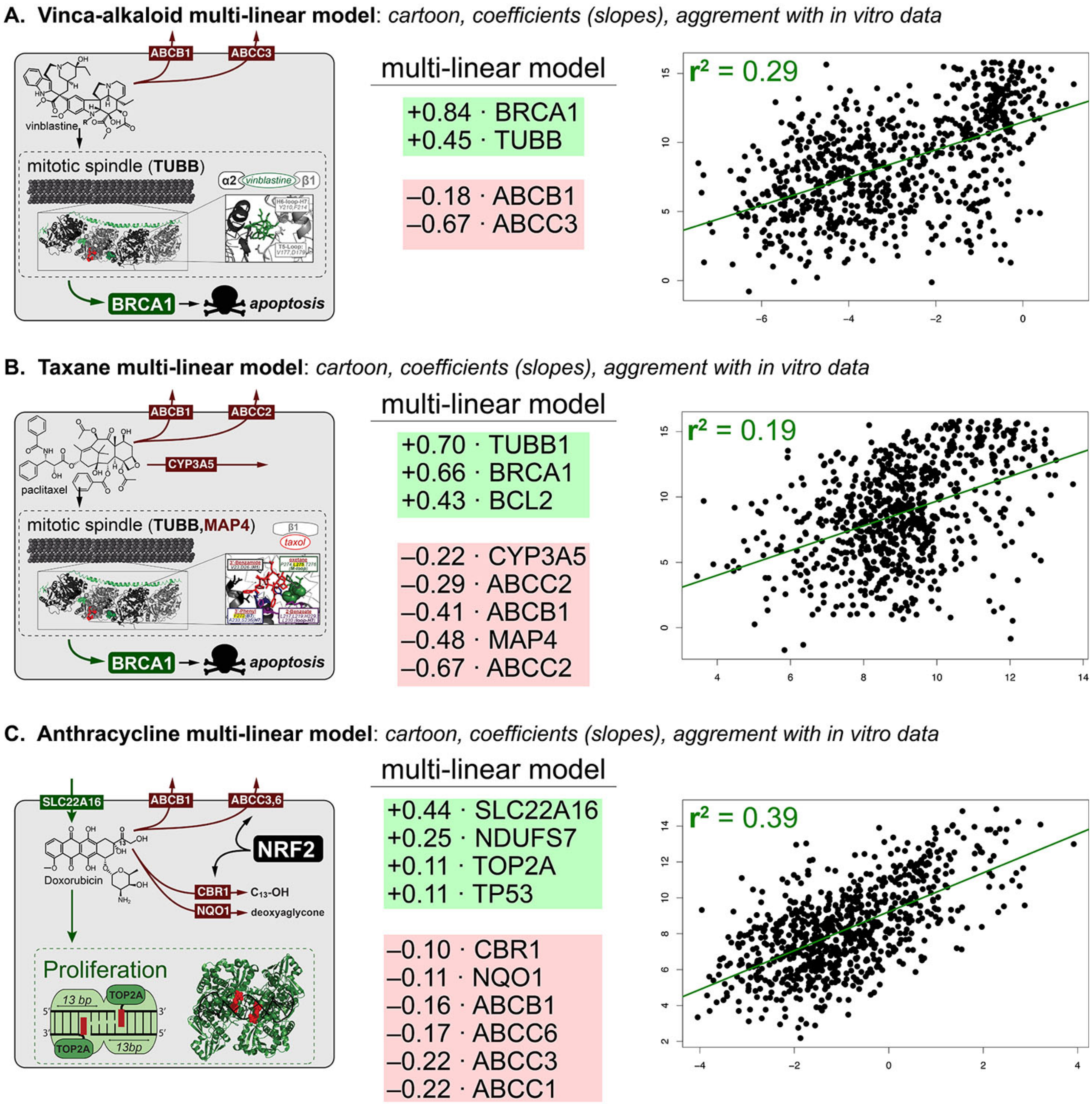

Figure S11 Multivariate Analyses of Top Biomarkers Independent Predictive Power. Multiple-linear regressions were performed on all causal biomarkers to deconvolute the individual contributions to chemotherapy sensitivity **(A)** Vinblastine multiple-linear model **(B)** Taxol multiple-linear model **(C)** Doxorubicin multiple-linear model

### Derivation of Chemoresistance EC50 Model

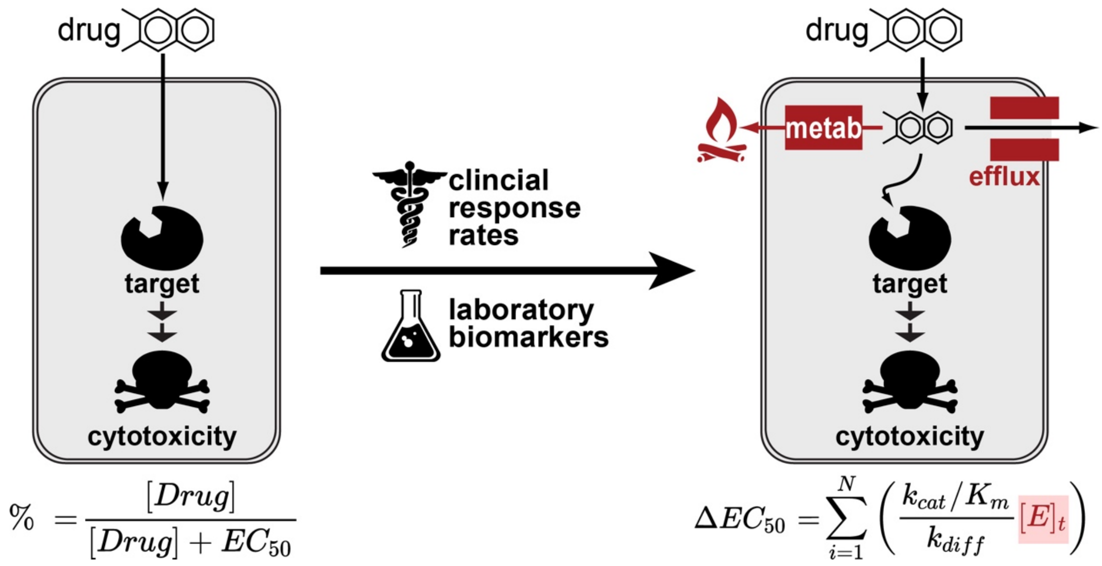

Figure S11 Overview of Classical (Hill) and New Model for Drug Potency (EC50). Classical Pharmacology theory assumes that drug potency (EC50) is a constant defining by the binding affinity of a drug for its target. We found that dose-response metrics (EC50/AUC) correlate better with expression of drug-metabolism and transport enzymes. This observation led to a new linear model that modifies the classical framework to better explain cell-type and tissue-variability in drug potency

#### Classical Pharmacology Theory (EC50 = Kd)

**Classical Pharmacology Theory.** Modern pharmacodynamics arose in the early 20th century with *receptor theory* that defined a drug effects as mediated by the physical binding of a drug (D) to a biological target (T) to form a complex (DT).[257] This theory was first quantified in 1910 by the Hill equation which defined the concentration of drug-target complex ([DT]) as a function of the target concentration ($[T]_t$), drug concentration ([*Drug*]) and a binding constant which quantified the strength of the drug-target interaction ($K_d$):[258,259]

$$[DT] = [T]_t \frac{[Drug]}{[Drug] + K_d} \quad \textbf{(1)}$$

The fractional term in equation 1 defines the fraction of drug-bound target which is assumed to be proportional to drug response (equation 2). The halfway point between 0% to 100% target binding is described by a specific drug concentration called the $EC_{50}$ or the Effective Concentration eliciting a 50% response:[257]

$$\%\, Response = \frac{[Drug]}{[Drug] + EC_{50}} \quad \textbf{(2)}$$

In 1913, Michaelis and Menten extended this framework to enzymatic processes, defining – for example – the rate at which a liver enzyme metabolizes a drug as proportional to fractional enzyme-binding, enzyme concentration ($[E]_t$) and the turnover number of the enzyme ($k_{cat}$).[260,261]

$$rate = k_{cat}[E]_t \frac{[Drug]}{[Drug] + K_m} \quad \textbf{(3)}$$

Since 1910, equations 1-2 have become the conceptual foundation for rational drug design which focuses on improving the potency of a drug ($EC_{50}$) by optimizing the drug's affinity for its target ($K_d$).[257] In parallel, equation 3 has proven foundational to the fields of biochemistry[262] and pharmacokinetics.[263] As detailed below, we derive a new model for drug-resistance which combines equations 1-3 to better predict cancer drug response.

**Limitations of Receptor Theory for Personalized Medicine.** Today, personalized cancer therapies are based on a 1-drug/1-gene paradigm derived from equation 1. Typically, a drug's target is measured (e.g. $[T]_t$) and used as a biomarker to predict drug response in individuals.[264,265] This approach has been critical to the success of targeted therapies, improving overall response rates (ORR) from 4% to 30% on average.[266,267]

While robust, this target-centric approach is clinically incomplete due to modest response rates.[264] In addition, this model is scientifically incomplete as dozens of non-target biomarkers have been discovered including drug transport and metabolism enzymes. These enzymes can affect a multi-drug resistance (MDR) phenotype and yet are not accounted for in standard pharmacodynamic theory (equation 1-2). Critically, pharmacokinetic theory provides a handle to model drug-transport/metabolism based on equation 3.

**Need for Personalized Cytotoxic Chemotherapies.** The limitations of this 1-drug/1-gene paradigm are particularly pronounced for cytotoxic chemotherapies (CC), which remain the backbone of metastatic cancer therapy[268,269] and are often combined with immunotherapies.[270] Despite decades of study, no target-based CC diagnostics have shown clinical utility.[265] As a result, CC regimens typically are applied based on cancer type (tissue-of-origin), which clinical guidelines have highlighted as insufficient.[271] Recently, several investigators have suggested that personalizing CC could represent a major paradigm shift[272] effecting the greatest clinical impact at the cheapest price.[268,269]

**Prioritizing N-biomarkers Required Clinical Validation:** Strikingly, hundreds of non-target CC-biomarkers have been identified in the laboratory, but none have translated to clinical practice.[265] For example, for non-small-cell lung cancer (NSCLC), only 13/80 predictive biomarkers have been investigated clinically and only 4 have shown any diagnostic potential.[269] Of these four, the drug-efflux pump ABCB1 is one of the most extensively studied yet only appears to explain ~15% of anthracycline resistance.[273] An emerging consensus is that no single biomarker is sufficient and multiple biomarkers will be needed to build robust CC diagnostics.[264,269,273] However, the lack of clinical evaluation has made it difficult to compare, prioritize and combine these biomarkers.[269,274]

Ideally, clinical validation should be driven by hypothesis-based clinical trials, but these are rare due to high expense and the limited availability of patient cohorts.[265] Retrospective studies on pooled clinical trials are becoming more common and it is this strategy that we use to assess laboratory-derived biomarkers of cytotoxic chemotherapy efficacy.[275,276]

#### New Chemoresistance Model Derivation

**Manuscript Summary:** A secondary goal of our single-agent database is two to evaluate the clinical validity of ~200 laboratory biomarkers for 34 of the top CC's.[277] Serendipitously, this evaluation led to the derivation of a pharmacodynamic model that reframes CC efficacy as a N-gene phenotype by combining equations 1-3. This model provides a molecular explanation for the selectivity of most clinical therapies as restricted to low drug-efflux and low drug-metabolism cancers.

**Mathematical models reconcile experimental and clinical literature**. Based on the superior predictive power of drug-metabolism and transport enzymes to predicting clinical and in vitro response we sought to mathematically define, from first principles, the contributions of these enzymes to standard pharmacodynamic theory (Equation 1-2). First, we assumed that the effect of these enzymes(E) occurs upstream to drug target binding where they primarily reduce the intracellular concentration of drug ($[drug]_{intra}$) within the cell (Figure 4E). Next, we assumed that this process could be modeled as parallel independent Michaelis-Menten-fluxes using equation 3:[262,278]

$$\boldsymbol{efflux} = \sum_{i=1}^{N} k_{cat}[E]_t \frac{[drug]_{intra}}{[drug]_{intra} + K_m} \qquad \textbf{4}$$

As the physiological role of most of these enzymes is hepatic and renal clearance, clinical regimens are designed to operate under 1st order conditions ([drug] < $K_m$) to prevent overdose.[263] As such, under physiological conditions, we can assume 1st order kinetics which simplifies equation 4 to:

$$\boldsymbol{efflux} = \sum_{i=1}^{N} \frac{k_{cat}[E]_t}{K_m} [drug]_{intra} \qquad \textbf{5}$$

After a certain amount of time (often minutes *in vitro*[279]), a steady state will be reached when the import and export rates of drug will be balanced:

$$influx = \boldsymbol{efflux} \qquad \textbf{6}$$

Assuming drug influx is primarily a diffusion limited process,[280] we can assign a first order rate constant ($k_{diff}$) and overall rate dependent on the extracellular concentration ($[drug]_{extra}$):

$$k_{diff}[drug]_{extra} = \sum_{i=1}^{N} \frac{k_{cat}[E]_t}{K_m}[drug]_{intra} \qquad \textbf{7}$$

Rearranging equation 7, we obtain a ratio of the extracellular and intracellular concentrations of a drug as a function of enzyme expression and 2N+1 physical constants:

$$\frac{[drug]_{extra}}{[drug]_{intra}} = \frac{\sum_{i=1}^{N} \frac{k_{cat}[E]_t}{K_m}}{k_{diff}} \qquad \textbf{8}$$

As the $EC_{50}$ defined by current pharmacodynamic theory is a drug concentration we can define this ratio as a change in $EC_{50}$ ($\Delta EC_{50}$). In addition, rearranging the right side of equation 8 we obtain an expression for differential drug-sensitivity as a weighted sum of drug-metabolism and drug-efflux enzyme concentrations:

$$\Delta EC_{50} = \sum_{i=1}^{N} \left( \frac{k_{cat}/K_m}{k_{diff}} [E]_t \right) \qquad \textbf{9}$$

Equation 9 provides a physical explanation for the surprising predictive power of the sum of biomarker-procedure illustrated in Figure 2A. Critically, this equation is based on first principles and makes the same underlying assumptions as standard pharmacokinetic theory.[262] As a result, it is likely that this expression has a similar scope (and limitations) as the Michaelis-Menten equation which is widely used in the laboratory and the clinic.
